# Circuit Degeneracy Facilitates Robustness and Flexibility of Navigation Behavior in *C.elegans*

**DOI:** 10.1101/385468

**Authors:** Muneki Ikeda, Shunji Nakano, Andrew C. Giles, Wagner Steuer Costa, Alexander Gottschalk, Ikue Mori

## Abstract

Animal behaviors are robust and flexible. To elucidate how these two conflicting features of behavior are encoded in the nervous system, we analyzed the neural circuits generating a *C. elegans* thermotaxis behavior, in which animals migrate toward the past cultivation temperature (*T_c_*). We identified multiple circuits that are highly overlapping but individually regulate distinct behavioral components to achieve thermotaxis. When the regulation of a behavioral component is disrupted following single cell ablations, the other components compensate the deficit, enabling the animals to robustly migrate toward the *T_c_*. Depending on whether the environmental temperature surrounding the animals is above or below the *T_c_*, different circuits regulate the same behavioral components, mediating the flexible switch between migration up or down toward the *T_c_*. These context-dependencies within the overlapping sub-circuits reveal the implementation of degeneracy in the nervous system, providing a circuit-level basis for the robustness and flexibility of behavior.

## Introduction

Animal behaviors exhibit two conflicting features, robustness and flexibility. Animals robustly execute behavior despite external and internal perturbations (Macmillan, 2000; Maddox, 1994; Meyer et al., 1998), but flexibly behave within a variable environment owing to the adaptation to external and internal changes (Honma et al., 2003; Okamoto and Aizawa, 2013; Saper et al., 2002). These two features lead animals to better chance of survival and reproduction, and also provide animals with higher evolvability (Edelman and Gally, 2001; Whitacre and Bender, 2010). Reconciling robustness and flexibility is thus a fundamental aspect for every biological system (Heinl and Grabherr, 2017; Kitano, 2004; Meir et al., 2002). However, it remains unclear how biological systems, especially the nervous system, can generate robust and flexible outputs.

With a compact nervous system consisting of only 302 neurons, the free-living nematode *Caenorhabditis elegans* exhibits navigation behaviors that are robust and flexible. Behaviors with such features are well exemplified by thermotaxis and chemotaxis. *C. elegans* animals can memorize environmental stimuli such as temperature or ion concentration in association with their feeding state (Hedgecock and Russell, 1975; Kunitomo et al., 2013). When placed on the region in a thermal gradient, where the temperature is higher than that of the past environment, animals migrate down the gradient toward the past cultivation temperature (T_c_). The migration is robust in a variety of thermal environments (Jurado et al., 2010; Ramot et al., 2008) and with deficiencies in the nervous system (Beverly et al., 2011; Luo et al., 2014a). By contrast, when placed on the region with a temperature lower than the *T_c_*, animals flexibly switch from migration down to up the gradient (Hedgecock and Russell, 1975; Ito et al., 2006). Also in chemotaxis, animals show both the robust execution of migrations (Iino and Yoshida, 2009; Luo et al., 2014b; Wang et al., 2017) and the flexible switching between migrations up and down salt gradients (Klein et al., 2017; Ohno et al., 2014). Nevertheless, how robust and flexible migration in thermotaxis and chemotaxis is achieved remains poorly understood.

A growing body of evidence suggests that neurons of *C. elegans* are multifunctional (Li et al., 2014; M. Liu et al., 2018), allowing a single neuron to contribute to multiple aspects of information processing within one or more circuits. By contrast, multiple distinct neurons often contribute to similar information processing (Beverly et al., 2011; Koo et al., 2011; Trojanowski et al., 2014). These one-to-many and many-to-one mappings in the nervous system, which are observed over many different animal species (Jankowska, 2001; Leonardo, 2005; Schiller, 1996; Shih et al., 2015), can be theoretically considered within the framework of *degeneracy* (Edelman and Gally, 2001; Tononi et al., 1999). Degeneracy refers to conditions where a system is conceptually composed of multiple subsystems whose components are partially shared. In this form, subsystems can coordinately produce the same output in some cases and produce different outputs from each other in other cases. Degeneracy is proposed to be one of a few strategies for a system to possess robustness and flexibility together (Wagner, 2005; Whitacre, 2010). Recently, two examples of *C. elegans* neurons with characteristics that suggest the implementation of degeneracy have been reported. 1) Depending on the temperature range during thermotaxis behavior, two thermosensory neurons, either AFD and AWC or AFD and ASI, are shown to be responsible for migration down a thermal gradient toward the *T_c_* (Beverly et al., 2011). 2) Pharyngeal interneuron I1 can excite and inhibit the pumping rate via two different pharyngeal motor neurons, MC and M2, independently (Trojanowski et al., 2014). However, it remained unknown whether and how neural circuits, working together as a network system, implement degeneracy and how robustness and flexibility emerge from such circuit systems as features of behavior.

Here, we addressed these questions by analyzing the neural circuits generating *C. elegans* thermotaxis. By combining high-throughput behavioral analysis and comprehensive cell ablations, we identified sub-circuits that regulate behavioral components, such as turns, reversals, and curves. These sub-circuits, required for the regulation of individual behavioral components, were distinct but highly overlapping. In a shared interneuron among sub-circuits, the regulation of different behavioral components was encoded in different ranges of neural activity according to the animals’ moving direction relative to the *T_c_*. We further found that when a deficiency in a sub-circuit was created, the behavioral components mediated by other sub-circuits compensated the defect, leading to the robust migration toward the *T_c_*. Depending on whether the animals are above or below the *T_c_*, similar but different sub-circuits generated opposing outputs in the same behavioral components, leading to the flexible switching between migrations up and down thermal gradients. Thus, our results demonstrate the implementation of degeneracy in the nervous system, identify a neural basis of circuit degeneracy, and show that circuit-level degeneracy ensures animals to execute robust and flexible behaviors.

## Results

### Migration toward the Cultivation Temperature Is Driven by the Flexible Regulations of Multiple Behavioral Components

*C. elegans* animals are known to navigate using a series of stereotyped movements, designated behavioral components (Croll, 1975; Iino and Yoshida, 2009; Pierce-Shimomura et al., 1999). We first attempted to extract the behavioral components during thermotaxis by employing a Multi-Worm Tracker (MWT) (Swierczek et al., 2011). MWT simultaneously captured the positions and postures of approximately 120 animals (Figure 1A), and these data were further analyzed by a custom-built MATLAB script to detect the behavioral components (see Materials and methods).

**Figure 1.**
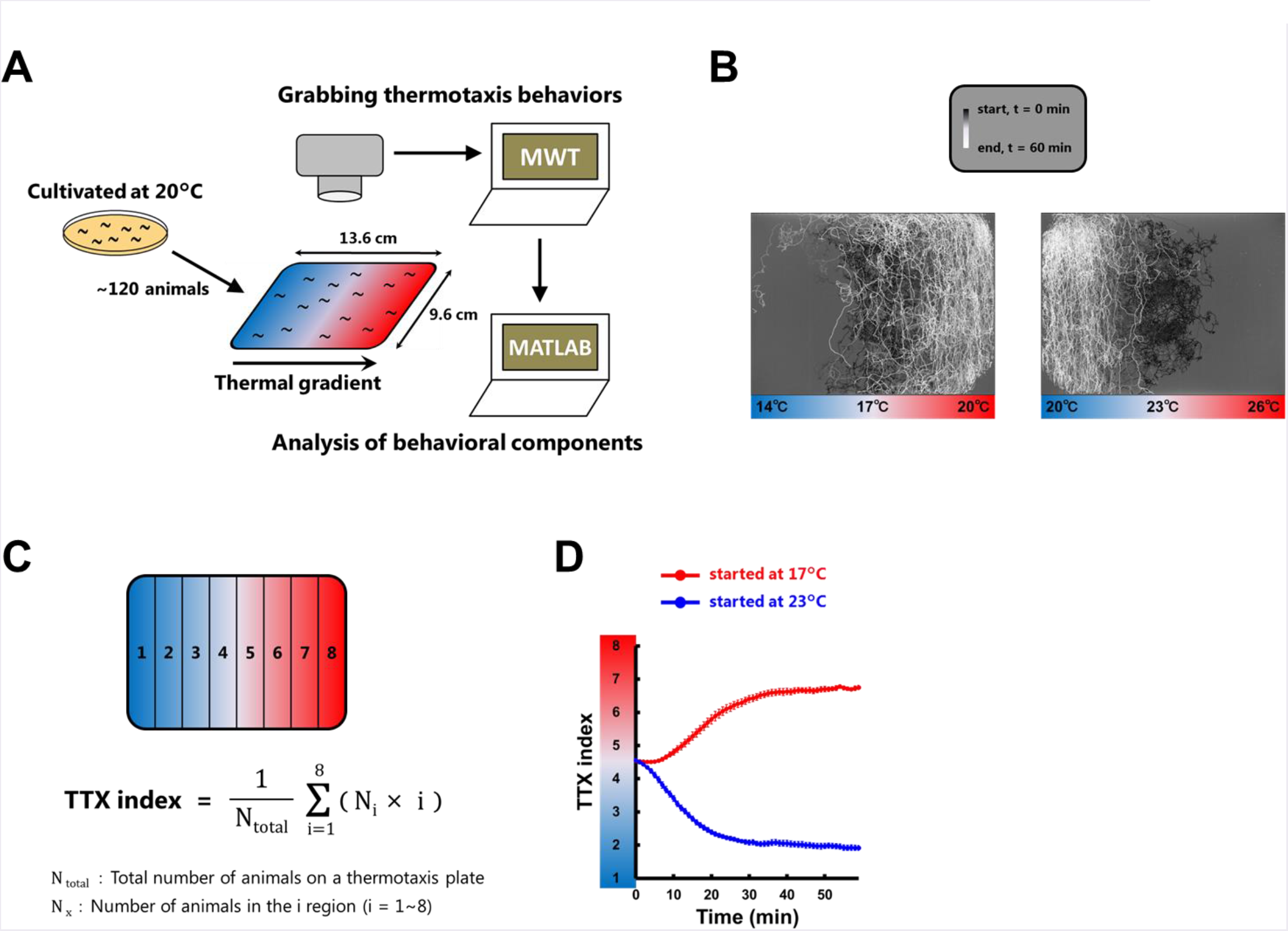
Thermotaxis behavior is accomplished within 30 minutes. (A) A Multi-Worm Tracker (MWT) system for the extraction of behavioral components during thermotaxis behavior. The thermotaxis assays were performed as previously reported (Ito et al., 2006). The positions and postures of animals were captured by the MWT system and then analyzed by custom-built MATLAB scripts. (B) Animals cultivated at 20°C were placed on a TTX plate with a thermal gradient with 17°C (left panel) or 23°C (right panel) at the center. Shown here are the representative trajectories of approximately 120 animals that were recorded by MWT. The time from the start of the assays are represented in gray scale. (C) Formula for the TTX index. The number of animals in each fraction (1-8) was scored in every one minute, and the TTX indices were calculated using the equations as described. (D) The time course of TTX indices in the *T<T_c_* condition (red line) and in the *T>T_c_* condition (blue line). The migrations toward the *T_c_* were almost accomplished within 30 minutes. (n = 12).

For the thermotaxis assays, we set cultivation temperature (T_c_) as 20°C and the temperature of the center in the assay plate as either 17°C or 23°C (Figure 1B). Animals were placed at the center of the plate, and then we evaluated the animals’ migrations by calculating thermotaxis index (TTX index), according to the equation shown in Figure 1C. TTX index is 1 when all the animals are in the coldest fraction of the plate and 8 when all the animals are in the warmest fraction. Consistent with our previous report (Ito et al., 2006), the animals reached their *T_c_* within approximately 30 minutes in two conditions (Figure 1D and Movie S1), plate centered at 17°C or 23°C. In this study, we thus focused on the first 30 minutes from the start of the assays. To analyze the behaviors during the migrations toward the *T_c_*, we analyzed the animals that were distributed in the center four fractions of the assay plate; 15.5–18.5°C for the *T<T_c_* condition and 21.5-24.5°C for the *T>T_c_* condition (Figure 2A). Behaviors were essentially classified into three behavioral components: turns, reversals, and curves (Figure 2B). Turns were further classified into omega turns and shallow turns (Kim et al., 2011; Schild and Glauser, 2013), and reversals were further classified into reversals and reversal turns (Croll, 1975; Pierce-Shimomura et al., 1999; Salvador et al., 2014).

**Figure 2.**
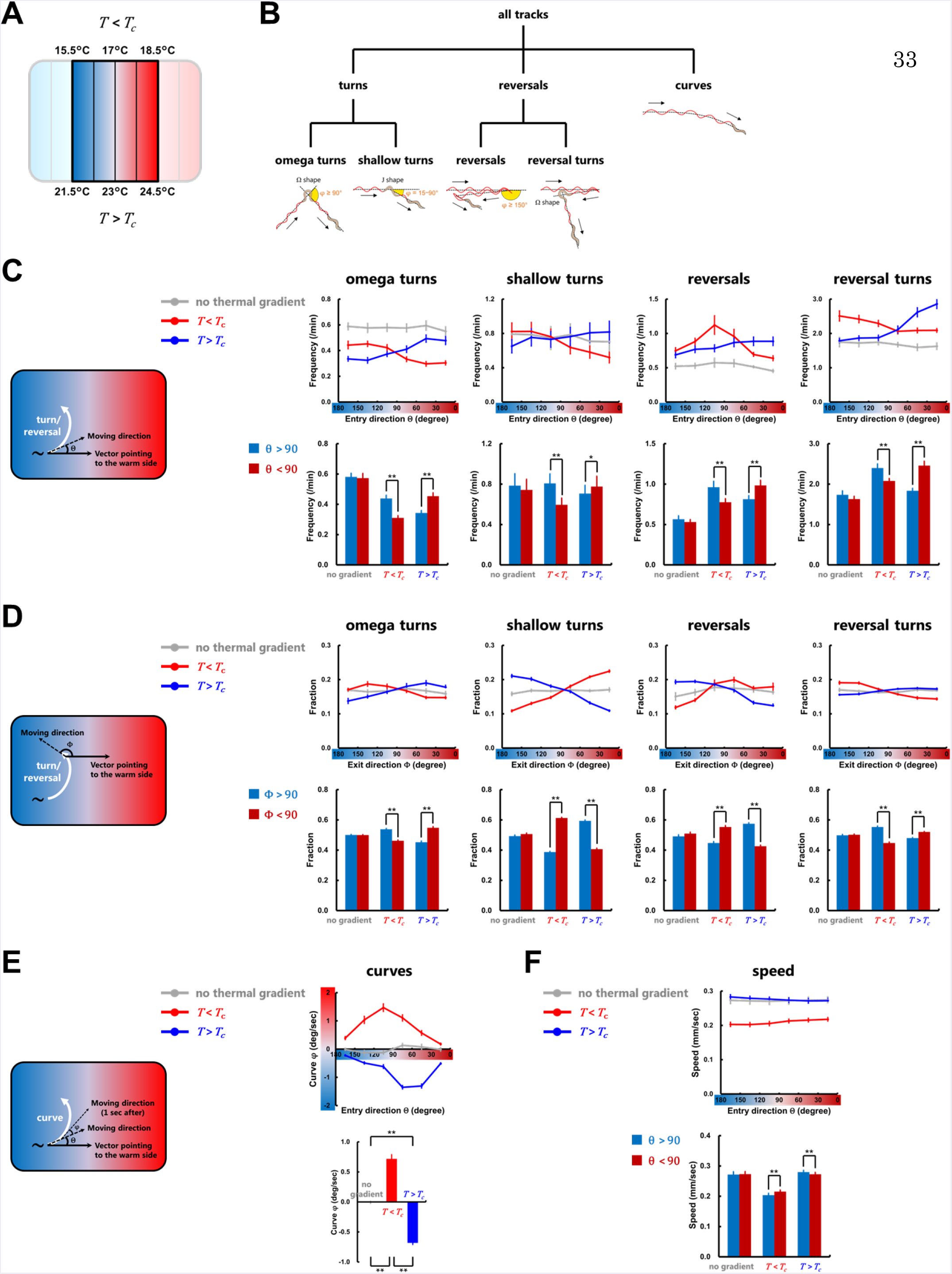
Behavioral components are flexibly regulated during thermotaxis. (A) Temperature range within which the behavioral components were analyzed. Animals in the center four fractions of the assay plate were analyzed (Figure 1B). (B) Classification and definition of *C. elegans* behavioral components used in this study. Turns, reversals, and curves were classified as previously proposed (Kim et al., 2011; Pierce-Shimomura et al., 1999; Salvador et al., 2014; Schild and Glauser, 2013). (C) Upper panels are frequency plots of the turns and the reversals representing the average as a function of the entry direction θ, the angle between the moving direction before these behavioral events and the vector pointing to the warm side of the thermal gradient. Lower panels show comparisons of the frequency of each behavioral event. Deep red columns indicate the frequencies of the events while the animals are moving up the thermal gradient (θ<90), and deep blue columns the frequencies while moving down the thermal gradient (θ>90). (D) Upper panels are fraction plots of the exit direction (Φ) after the turns and the reversals. Φ is the angle between the moving direction after these behavioral events and the vector pointing to the warm side of the thermal gradient. Lower panels show comparisons of the fraction of the exit direction of each behavioral event. Deep red columns indicate the exit directions of the events toward the warm side of the thermal gradient (Φ<90), and deep blue columns the exit directions toward the cold side (Φ>90). (E) The biases (φ) of curves were calculated for each frame and averaged as a function of the moving direction θ. φ is the angle between the past moving direction (1 sec before) and the current moving direction. φ is defined as positive if biased toward higher temperature and negative if biased toward lower temperature. (F) Upper panel is speed plots representing the averages as a function of the entry direction θ. Lower panel shows comparisons of the speeds. Deep red columns indicate the speeds while the animals are moving up the thermal gradient (θ<90), and deep blue columns the speeds while moving down the thermal gradient (θ>90). In (C-F), gray lines correspond to experiments without the thermal gradient (20°C constant), red lines correspond to experiments in the *T<T_c_* condition, and blue lines correspond to experiments in the *T>T_c_* condition. n ≥ 6. Error bars indicate SEM. Paired Student’s t-test (C, D, F); one-way ANOVA followed by a Tukey-Kramer post hoc multiple comparisons test (E). **p < 0.01, *p < 0.05.

Our analyses show that the behavioral components were oppositely biased depending on whether the animals were moving below or above the *T_c_ (T<T_c_* or *T>T_c_* conditions, respectively) (Figures 2C-E). The frequencies of turns and reversals were measured as a function of the entry directions θ, where θ is the difference between the moving direction right before these behavioral events and the vector pointing to the warm side of the thermal gradient (Figure 2C). In the *T<T_c_* condition, the frequencies of turns and reversals were higher when the animals were moving down the thermal gradient (θ>90) than when the animals were moving up the thermal gradient (θ<90), whereas in the *T>T_c_* condition, the frequencies were higher when the animals were moving up the thermal gradient (θ<90) than moving down the thermal gradient (θ>90) (Figure 2C). We also measured the fraction of the exit direction Φ after the turns and the reversals, where Φ is the angle between the direction right after these behavioral events and the vector pointing to the warm side of the thermal gradient (Figure 2D). The exit directions of shallow turns and reversals were biased toward the *T_c_* both in the *T<T_c_* and *T>T_c_* conditions; biased toward the warm side (Φ<90) in the *T<T_c_* condition whereas biased toward the cold side (Φ>90) in the *T>T_c_* condition (Figure 2D).

We also analyzed two components associated with forward movement: the curve direction and the locomotion speed. Curve direction was measured as the angle φ, where φ was the angle between the past moving direction and the current moving direction (Figure 2E). Similar to the exit directions of shallow turns and reversals, curve direction was biased toward the warm side (φ>0) in the *T<T_c_* condition and biased toward the cold side (φ<0) in the *T>T_c_* condition (Figure 2E). The locomotion speed also showed the opposite bias under the *T<T_c_* or *T>T_c_* conditions. In the *T<T_c_* condition, the locomotion speed was faster when the animals were moving up the thermal gradient (θ<90) than when the animals were moving down the thermal gradient (θ>90), whereas in the *T>T_c_* condition, the locomotion speed was faster when the animals were moving down the thermal gradient (θ>90) than moving up the thermal gradient (θ<90) (Figure 2F). It should be noted that in the *T>T_c_* condition, the biases of turns, reversal, and curves were observed in the time windows earlier than the biases in the *T<T_c_* condition (Figures S1A-S1E), which may enable the faster migration toward the *T_c_* when the animals are moving down toward the *T_c_* than moving up toward the *T_c_* (Ito et al., 2006; Luo et al., 2014a) (Figure 1D).

Taken together, our results suggest that sensory inputs of temperature increment and decrement are processed differently according to the relative position of the animals to the *T_c_*, in which the behavioral components were oppositely regulated in *T<T_c_* and *T>T_c_* conditions, thereby enabling the animals to migrate toward the *T_c_*.

### Different Sets of Behavioral Components Are Employed Depending on Different Temperature Environment Relative to the *T_c_*

Although the biases of behavioral components likely play essential roles in thermotaxis behavior (Figure 2), whether those biases are necessary and sufficient to drive the animals to migrate toward the *Tc* is unclear. To address this question, we conducted Monte Carlo simulations of animals’ migrations on the thermal gradient. In the simulation, we defined an animal’s state by its position in the assay plate (x, y) and the direction of its movement relative to the vector pointing to the warm side of the thermal gradient (θ) (Figure 3A). We updated the states of animals every second, according to the experimental data of the turning frequencies, the exit directions of turns and curves (Φ and φ), and speeds (v), as functions of θ and *T* versus *T_c_* (Figure S1A-S1F, see Materials and methods). Similar to the animals observed in thermotaxis assays, computer-simulated animals (sims) moved up and down the thermal gradient toward the *Tc* within 30 minutes (Figure 3B and Movie S2), suggesting that the behavioral biases shown in Figure 2 are sufficient for the animals to reach the *T_c_*.

**Figure 3.**
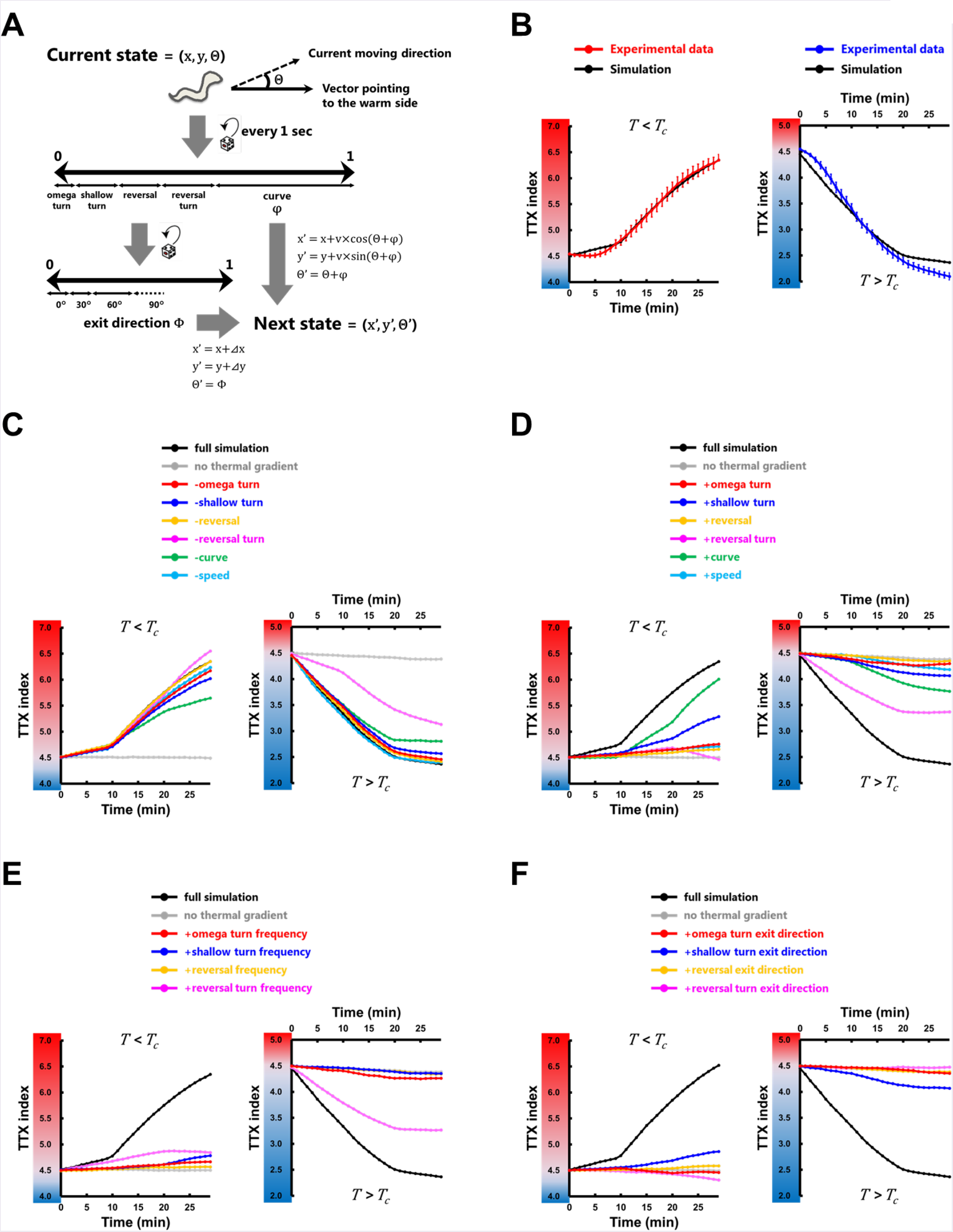
Behavioral components are employed differently depending on different temperature environment relative to the *T_c_*. (A) Schematic structure of the thermotaxis behavior simulation. Animal’s state was defined by its position (x, y) and moving direction relative to the vector pointing to the warm side (θ). We updated the states of the animal every second according to the experimentally observed data: the frequencies and the exit directions (Φ) of the turns and the reversals, the biases of the curves (φ), and the speeds (v) (Figures 2C-2F, S1A-S1E), all of which were applied as functions of θ and *T* versus *T_c_*. The displacements during the individual turns *(Ax*, Ay) were also employed when updating the states of the animals. (B) The time course of TTX indices in the simulations (black lines) and that obtained from experimental data (colored lines). In the simulations, we iterated assays for 100 times, each with 100 animals, and the TTX indices were averaged within the assays. (C) The time course of TTX indices in the simulations in which the data of the individual behavioral component determined by the experiment with the thermal gradient was replaced with the data of the corresponding component without the gradient. (D-F) The time course of TTX indices in the simulations in which the data of the individual behavioral component without the thermal gradient was replaced with the data of the corresponding component with the gradient. (D) Data of both frequencies and exit directions were replaced. (E) Data of frequencies alone was replaced. (F) Data of exit directions alone was replaced. In (C–F), black lines correspond to the simulation in which all the data of wild-type animals determined by the experiment with the thermal gradient were used, and gray lines correspond to the simulation in which all the data of wild-type animals without the gradient were used. The other colored lines correspond to the simulation with the replacements of the individual behavioral component: the omega turn (red lines), the shallow turn (blue lines), the reversal (yellow lines), the reversal turn (magenta lines), the curve (green lines), and the speed (light blue lines).

The simulation was also executed to examine the contributions of the individual behavioral components. First, we simulated the situations in which the sims could not use one of the behavioral components (see Materials and methods). In the *T<T_c_* condition, the removal of the curves impaired the increase of the TTX index most severely. In the *T>T_c_* condition, the removal of the reversal turns impaired the decrease of the TTX index most severely (Figure 3C). Second, we performed the simulations in which the sims used only one of the behavioral components. In the *T<T_c_* condition, the sims using only the curves showed the most dramatic increase of the TTX index (Figure 3D), whereas the sims using only the reversal turns showed little increment of the index. By contrast, in the *T>T_c_* condition, the reversal turns were most effective in decreasing the TTX index, and the curves also significantly caused a decrement of the TTX index. These analyses suggest that different sets of behavioral components contribute to the migration of the animals toward the *T_c_* depending on the context (T versus T_c_).

We further assessed separately the contributions of the biases in the frequencies and the exit directions. The simulations in which the sims used only the frequency of reversal turns showed the increase of the TTX index (Figure 3E), whereas the sims allowed to use only the exit direction did not show any increment (Figure 3F). By contrast, the simulations in which the sims using only the exit direction of shallow turns showed an increase of the index (Figure 3F), whereas the sims used only the frequency showed little increment (Figure 3E). These results showed that the bias in the frequency of reversal turns contributes to the migration of the animals toward the *T_c_*, whereas the bias in the exit direction of shallow turns contributes to the migration.

Although these results are consistent with the possibility that the difference of the temperature environment relative to the *Tc* is responsible for the different employment of the behavioral components (Figures 3C and 3D), it is also possible that the difference of absolute temperature may be responsible, since the animals are migrating in different temperature ranges (Figure 2A). Indeed, temperature itself is known to affect the turning frequencies and the speed of animals (Ryu and Samuel, 2002) (Figure S1G). To exclude the effect of absolute temperature, we next fixed the temperature of the center in the assay plate as 20°C and set the *T_c_* as 17°C or 23°C (Figures S2A and S2B). Also in these two conditions, our thermotaxis simulation reproduced the experimental data (Figure S2H). The simulations that disabled or allowed different individual behavioral components showed that 23°C cultivated animals moved up the thermal gradient toward the *T_c_* by employing mainly the curves, whereas 17°C cultivated animals moved down the thermal gradient toward the *Tc* by employing mainly the reversal turns and the curves (Figures S2I and S2J). These results are similar to the results when *T_c_* was constant and the center temperature varied (Figure 3), suggesting that context, not absolute temperature, is the important factor controlling impact of various behavioral components. Taken together, our results suggest that the animals switch the thermotactic behavioral strategies depending on the context of the temperature inputs, below or above the *T_c_*.

### Overlapping but Distinct Neural Circuits Are Recruited for the Context-dependent Regulation of Individual Behavioral Components

To investigate how the biases of the behavioral components are encoded in neural circuits during thermotaxis behavior, we attempted to identify the neurons that regulate the behavioral components. Ablations of individual neurons (Figure 4A) were performed by expressing reconstituted caspases (Chelur and Chalfie, 2007) or mito-miniSOG (Qi et al., 2012) (Table S1, see Materials and methods).We ablated individual neurons such as thermosensory neurons AFD, AWC and ASI (Beverly et al., 2011; Biron et al., 2008; Kuhara et al., 2008; Mori and Ohshima, 1995), locomotory command interneurons AVA and AVE and head motor neurons RMD, RME, SMB, and SMD that had been previously shown to regulate a backward locomotion or a steering behavior, respectively (Chalfie et al., 1985; Hart et al., 1995; Hendricks et al., 2012; Kocabas et al., 2012), and a series of interneurons that are predicted to be critical for mediating information transmission or integration (Donnelly et al., 2013; Kotera et al., 2016; Li et al., 2014; Ma and Shen, 2012). To evaluate how individual cell ablations affect the regulations of the behavioral components, we performed the thermotaxis simulations by using the experimental data of each behavioral component in the cell-ablated animals (Figures S5-S7). To quantify the performance of the simulations, we calculated migration index following the equation shown in Figure 4B. The difference between the TTX indices of cell-ablated animals and the indices of wild-type animals at a constant temperature was calculated every one minute and then summed up within 1-30 min. The value was divided, for normalization, by the summation of the difference between the indices of wild-type animals on the thermal gradient and the indices on the constant temperature. This index is 0 when biases of the behavioral components do not achieve any migration toward the *T_c_* and +1 (−1) when biases achieve the same migration as wild-type animals in the *T<T_c_ (T>T_c_)* condition.

**Figure 4.**
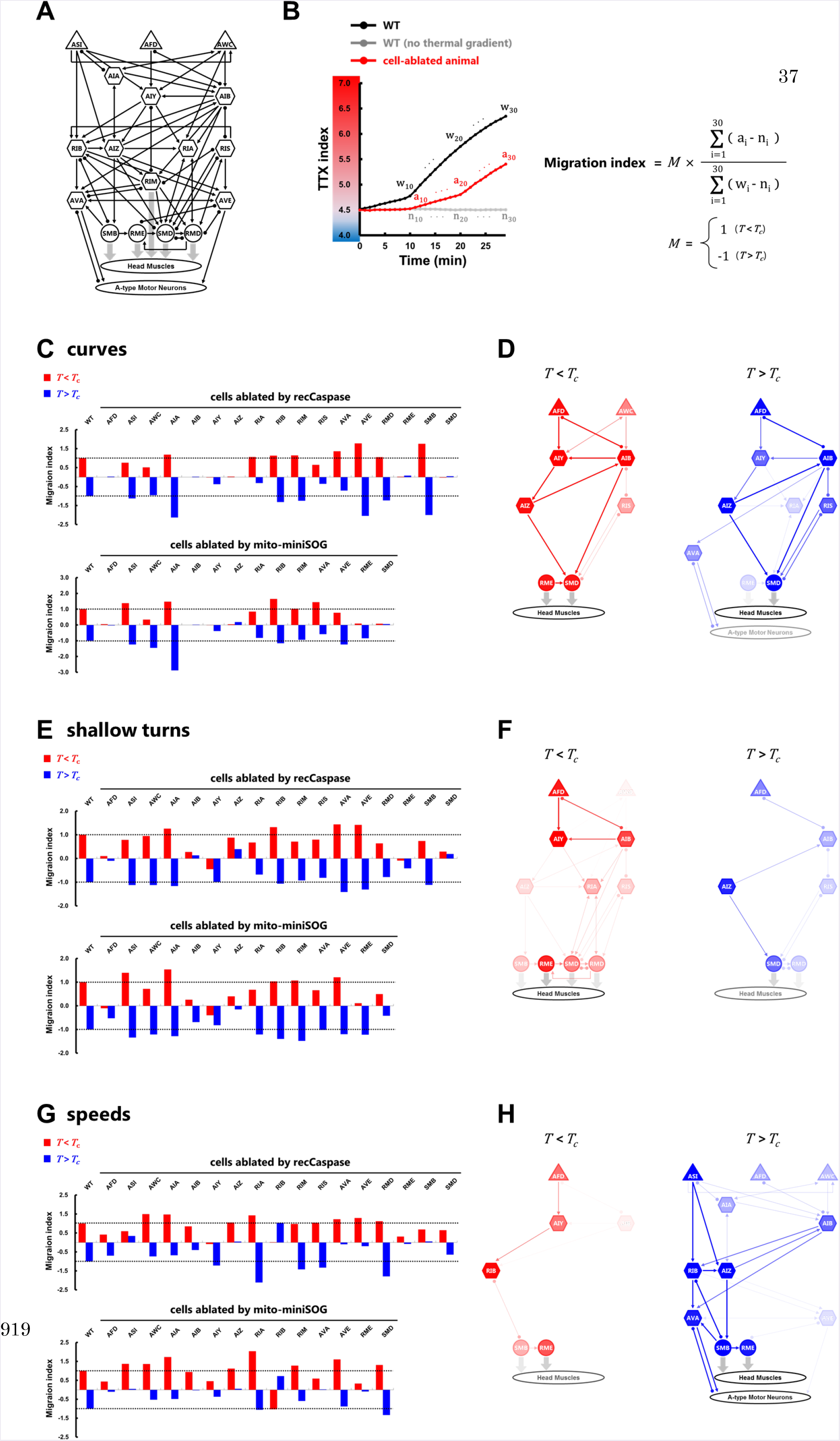
Overlapping but distinct neural circuits are recruited for the context-dependent regulation of the curves, the omega Turns, and the shallow turns. (A) Candidate neurons for cell-specific ablations, including thermosensory neurons (triangles), interneurons (hexagons), and head motor neurons (circles). Black thin arrows indicate chemical synapses, black undirected lines with round endings gap junctions, and gray thick arrows neuromuscular junctions. (B) Formula for the migration index. TTX indices from the simulation of cell-ablated animals (a_i_, red line) were compared with the indices from the simulation of wild-type (WT) animals without the thermal gradient (n_i_, grey line) in every minute, and the difference between them was summed up within 1-30 min. The value was normalized with the summation of the difference between the TTX indices from the simulation of WT with the thermal gradient (w_i_, black line) and the indices of WT without the gradient. (C, E, G) Migration indices of the curves, the shallow turns, and the speeds after cell-specific ablations by expressing reconstituted caspases (upper panels) and mito-miniSOG (lower panels). The indices in the *T<T_c_* condition are represented as red columns. The indices in the *T>T_c_* condition are represented as blue columns. Dashed lines show the indices of wild type animals (±1). (n ≥ 5). (D, F, H) Predicted neural circuits for regulating the curves, the shallow turns, and the speeds in the *T<T_c_* condition (red) and in the *T>T_c_* condition (blue). The thickness and color strength of each neuron represent the functional importance of the neuron predicted from the analysis and were determined as follows: For each neuron, the differences between the migration index of the wild-type animals and the index of the cell-ablated animals expressing reconstituted caspases or mito-miniSOG were calculated. The smaller difference from the two ablation strategies is used to determine the color strength, where the color strength of each neuron is proportional to this value. The color strength of each line is identical to the strength of the color of one of the two connected neurons with lower strength, and the thickness of each line is proportional to this color strength.

The ablations of AFD, AIB, AIZ, and SMD abolished the indices of the curves in both *T<T_c_* and *T>T_c_* conditions (Figure 4C), suggesting that these neurons play key roles in the regulation of the curves. Interestingly, the ablations of other neurons had different impacts on the indices of the curves under the *T<T_c_* or *T>T_c_* conditions. For example, the ablation of AWC impaired the indices only in the *T<T_c_* condition, whereas the ablations of RIA and AVA impaired the indices only in the *T>T_c_* condition. The index of the AIY-ablated animals was almost zero in the *T<T_c_* condition but had non-zero value in the *T>T_c_* condition. These results suggest that the different neurons are recruited to regulate the curves under the *T<T_c_* or *T>T_c_* conditions (Figure 4D). We applied the same analyses on the exit directions of the shallow turns (Figures 4E and S5C) and the speeds (Figures 4G and S5E). The indices of the shallow turns in the AIY-ablated animals were abolished (Figure 4E) in the *T<T_c_* condition but remained relatively normal in the *T>T_c_* condition. The indices of the speeds in the ASI-ablated animals and the AIZ-ablated animals were abolished in the *T>T_c_* condition but remained relatively normal in the *T<T_c_* condition (Figures 4G). These results suggest that different neural circuits are also recruited to regulate the shallow turns and the speeds under the *T<T_c_* or *T>T_c_* conditions (Figures 4F and 4H).

### Redundancy and Compensatory Interactions among Sub-circuits Ensure Robust Migration toward the *T_c_* under the Deficient Circuits

Unlike the case of the curve, the shallow turn, and the speed, none of the ablations of single sensory neuron abolished the migration index of the reversal turns (Figures 5A and S6A), which were employed mainly in the *T>T_c_* condition (Figures 3C-3E). To examine whether AFD and AWC sensory neurons, each of which shows small contribution to the indices (Figures 5A), redundantly regulate the reversal turns, we silenced AFD and AWC simultaneously. The indices of AFD-AWC double ablated animals were impaired more severely than those of the single ablated animals (Figures 5C and S6B). We further analyzed the reversal turns of *gcy-23 gcy-8 gcy-18 ceh-36* quadruple mutants, where *gcy-23 gcy-8 gcy-18* triple mutants and *ceh-36* mutants have been regarded as the AFD-deficient animals and the AWC-deficient animals, respectively (Inada et al., 2006; Lanjuin et al., 2003). Similar to the AFD-AWC double ablated animals, the index of *gcy-23 gcy-8 gcy-18 ceh-36* quadruple mutants was abolished and was lower than the indices of *gcy-23 gcy-8 gcy-18* triple mutants and *ceh-36* mutants (Figures 5C and S6B). The index of *eat-4* mutants, which encodes the vesicular glutamate transporter EAT-4 expressed in both AFD and AWC (Ohnishi et al.,2011), was also completely impaired (Figures 5D and S6C). This phenotype was partially rescued by AFD-specific expression or AWC-specific expression of a wild-type *eat-4* cDNA (Figures 5D and S6B). These results support the idea that temperature input from both AFD and AWC are required to fully achieve the regulation of the reversal turns and that input from only AFD or only AWC can be independently processed to regulate the reversal turns.

**Figure 5.**
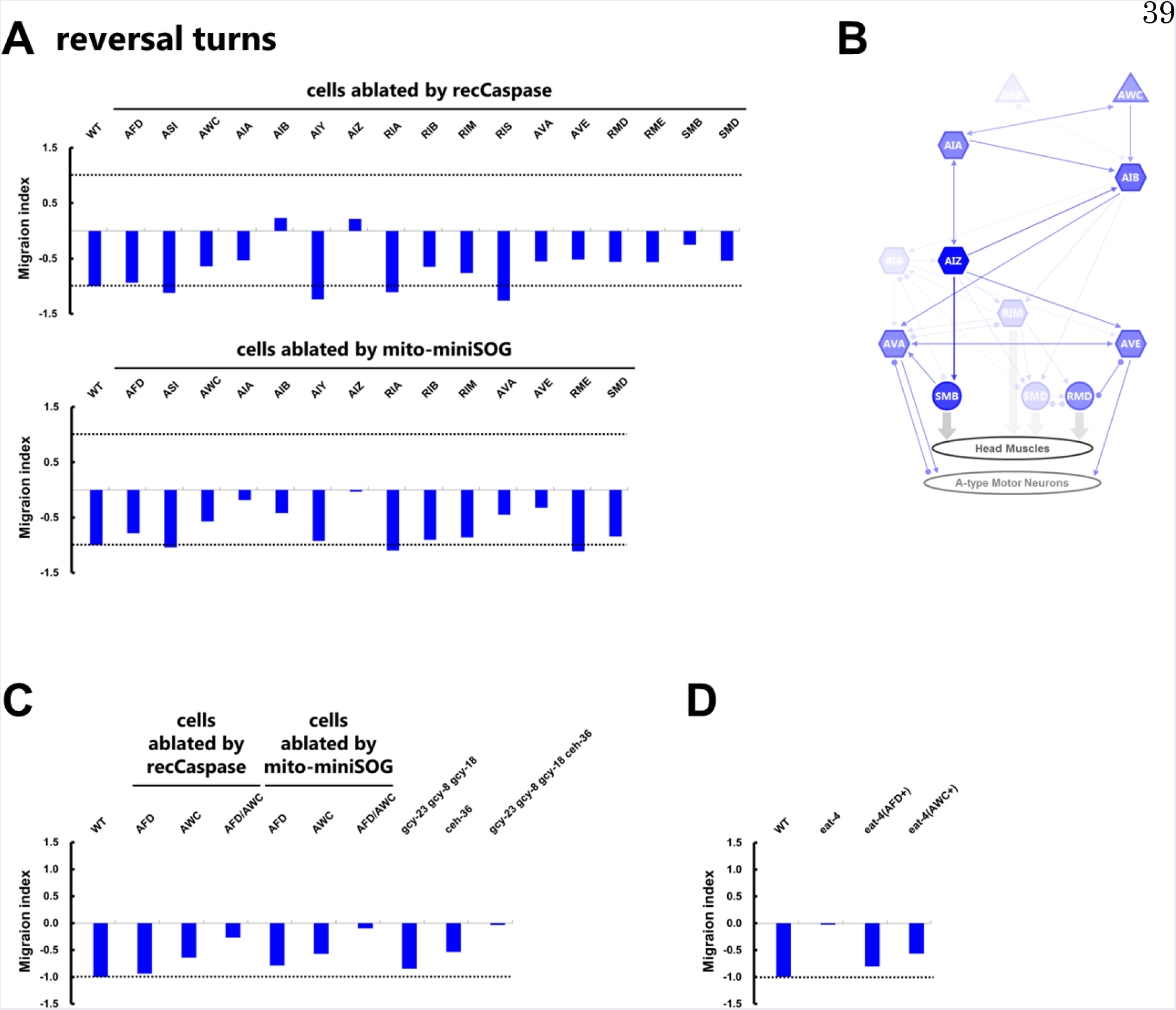
Redundancy between AFD and AWC sensory neurons enables robust regulation of the reversal turns. (A) Migration indices of the reversal turns after cell-specific ablations by expressing reconstituted caspases (upper panel) and mito-miniSOG (lower panel). (n ≥ 5). (B) Predicted neural circuits for regulating the reversal turns. (C) Migration indices of the reversal turns of wild-type animals, AFD-deficient animals, AWC-deficient animals, and AFD-AWC double deficient animals. (n ≥ 5). (D) Migration indices of the reversal turns of wild-type animals, *eat-4* mutants, *eat-4* mutants with an expression of *eat-4* cDNA in AFD, and *eat-4* mutants with an expression of *eat-4* cDNA in AWC. (n ≥ 6). In (A, C, D), dashed lines show the indices of wild type animals (−1).

We also noticed that the impairments in the index of a behavioral component sometimes accompany with the enhancement in the index of other behavioral components. The ablations of AIA, AVE, and SMB impaired the indices of the speeds and the reversal turns (Figures 4G and 5A), whereas the indices of the curves and the omega turns of these cell-ablated animals were larger than the indices of wild-type animals (Figures 4C and 6A). Also, RIS-ablated animals showed the impaired index of the curves and the shallow turns (Figures 4C and 4E) but showed the enhanced index of the reversal turns (Figure 5A). Since the TTX indices in the assays showed that RIS-ablated animals and AVE-ablated animals migrated toward the *T_c_* as successfully as wild-type animals (Figures 6D and 6E), the enhanced regulations of the behavioral components might help the migrations of the cell-ablated animals. Indeed, when we simulated the situations in which the reversal turn of RIS-ablated animals were replaced with those of the wild-type animals (Figure 6G), the decrement of the TTX index was partially prevented, suggesting that the enhanced regulation of the reversal turns helps the migration of RIS-ablated animals. The same kind of analyses on AVA-ablated animals also supported the compensatory interaction among the behavioral components (Figure 6H). It should be noted that none of the single-cell ablations, including the ablation of the sensory neurons and the motor neuron (Figures 6B and 6F), completely eliminated the migration toward the *T_c_*. These results suggest that deficiencies in the nervous system are compensated by the sub-circuits regulating the behavioral components to execute the robust migration toward the *T_c_*.

**Figure 6.**
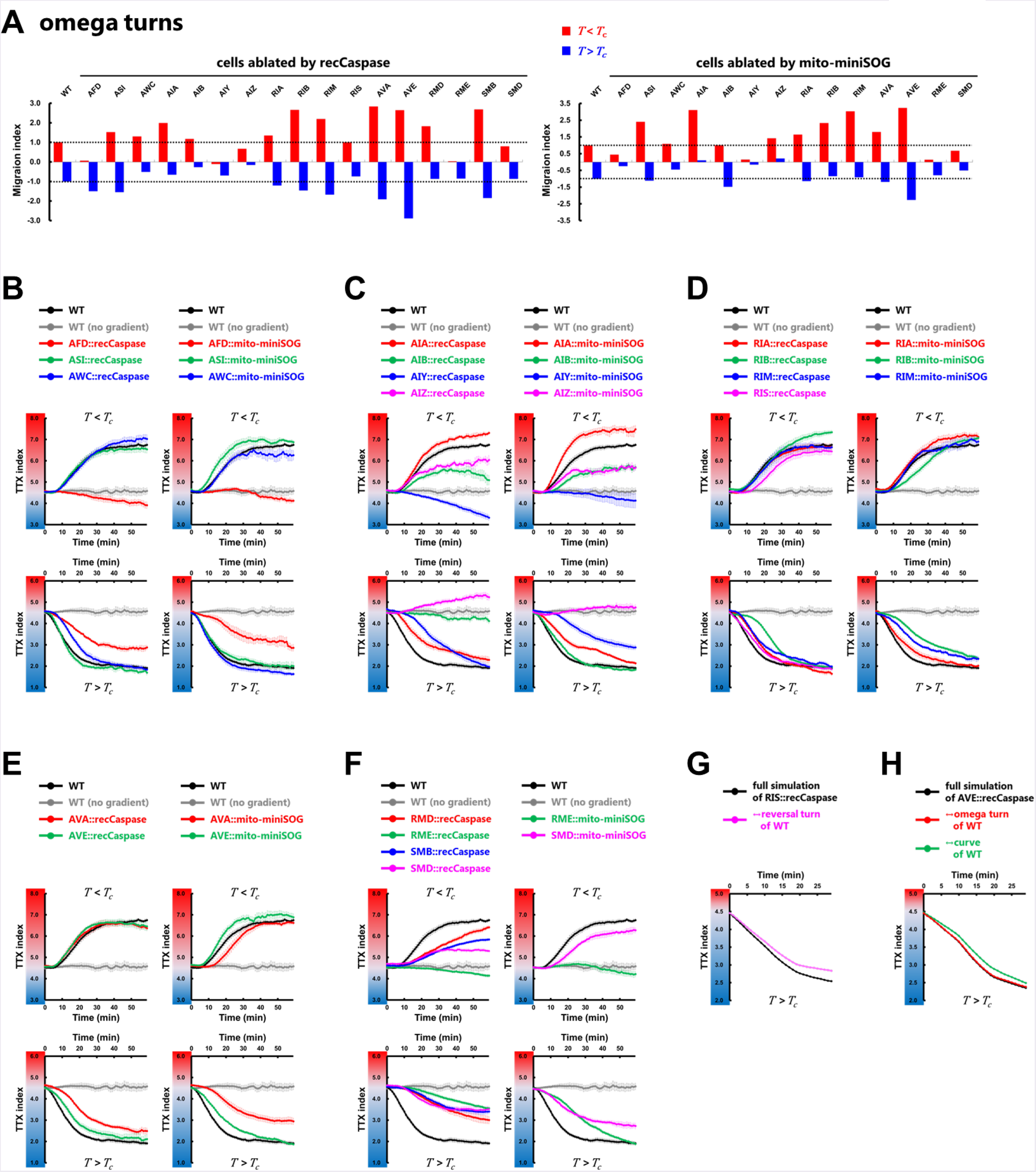
The migration toward the *Tc* is robustly executed under the deficiencies in the neural circuits. (A) Migration indices of the omega turns after cell-specific ablations by expressing reconstituted caspases (left panel) and mito-miniSOG (right panel). The indices in the *T<T_c_* condition are represented as red columns. The indices in the *T>T_c_* condition are represented as blue columns. Dashed lines show the indices of wild type animals. (n ≥ 5). (B-F) The time course of TTX indices in the *T<T_c_* condition (upper panels) and in the *T>T_c_* condition (lower panels) under the deficiency of thermosensory neurons (B), amphid interneurons (C), ring interneurons (D), ventral cord interneurons (E), and ring motor neurons (F). n ≥ 5. Error bars indicate SEM. (G) The time course of TTX indices in the simulations of RIS-ablated animals (black line) and in the simulations in which the data of the reversal turns of RIS-ablated animals was replaced with the data of wild-type animals (magenta line). (H) The time course of TTX indices in the simulations of AVE-ablated animals (black line) and in the simulations in which the data of the omega turns and the curves of AVE-ablated animals was replaced with the data of wild-type animals (red line and green line, respectively).

### Context-dependent Information Processing in a Single Interneuron Enables the Regulation of Multiple Behavioral Components

To understand how multiple behavioral components can be regulated in overlapping sub-circuits, we monitored the neural activity of AIB interneuron, which regulates both curves and the reversal turns in the *T>T_c_* condition (Figures 4D, 5B, and 7A). We 2_+_ performed Ca^2+^ imaging in freely moving animals using the single-worm tracking (SWT) system (Tsukada et al., 2016) (Figure 7B). Like in MWT, we observed the bias of both curve and reversal turn frequency using SWT (Figure 7C); when the animals were moving up the thermal gradient (θ<90 i.e. dT/dt>0), the bias of the curves was stronger and the frequency of reversal turns was higher. Thus, we assessed the relationship between the activity of AIB and the behavioral components under the dT/dt>0 and dT/dt<0 situations. Under the dT/dt>0 situation, the stronger bias of the curves was accompanied by the higher activity of AIB (Figure 7D), whereas there was no significant correlation between the reversal turn frequency and the AIB activity (Figure 7E). By contrast, under the dT/dt<0 situation, there was no significant correlation between the curve bias and the AIB activity, whereas lower frequency of reversal turns was accompanied by lower activity of AIB. We also checked the relationship between the activity of AIB and the differential of temperature input. As 2_+_ shown in the histogram (Figure 7F), the significant difference of the Ca signal was not observed under the dT/dt>0 or dT/dt<0 situations (Figure 7G). These results suggest that different ranges of the AIB activity regulates the different behavioral components under the different situations; low AIB activity suppresses the frequency of the reversal turns when the animals are moving up the thermal gradient, and high AIB activity promotes the curves when moving down the thermal gradient. On the other hand, AIA interneuron, which mainly regulates the reversal turns in the *T>T_c_* condition (Figures 5B and S8A), showed higher activity accompanied with lower frequency of reversal turns under both dT/dt<0 and dT/dt>0 situations (Figure S8G). In summary, the context-dependent regulation of behavioral components by different ranges of neural activity may enable one-to-many mappings between single neurons and multiple behavioral components, underlying the implementation of degeneracy.

**Figure 7.**
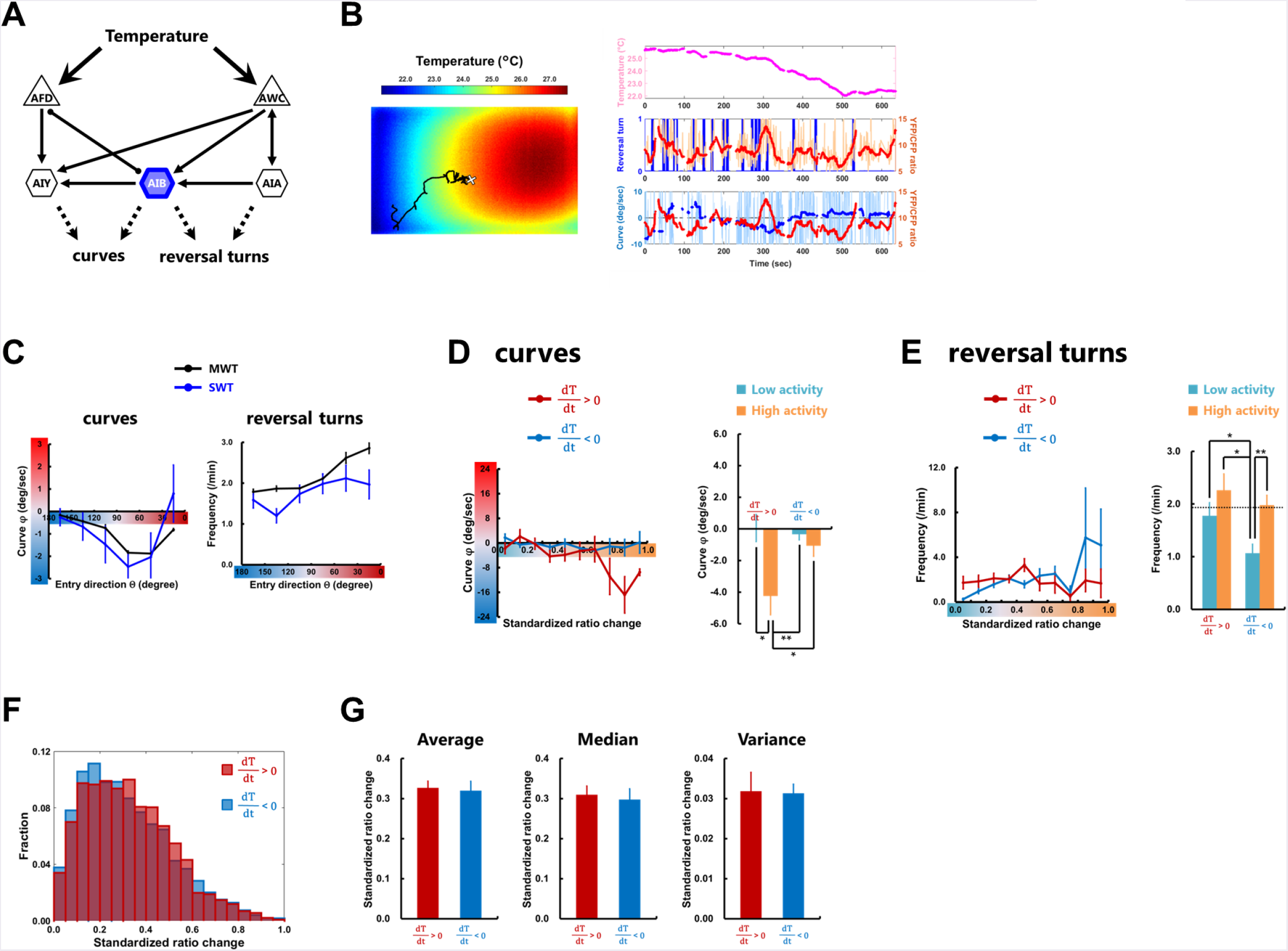
AIB neuron regulates the curves and the reversal turns in a context-dependent manner. (A) Representative sensory neurons and first-layer interneurons, which regulate the reversal turns or the curves. Thin arrows indicate chemical synapses and an undirected line with round endings indicates gap junction. (B) Representative thermography image (left panel) taken together with calibrating temperature measurements using a thermocouple sensor. Projection of a trajectory (black line) shows how the animal searches the thermal environment. The white + marks the starting point when recording starts. From this trajectory, the time course of the temperature changes is obtained (light magenta line in right upper panel), and the reversal turn (blue line in right middle panel) and the curve (light blue line in right lower panel) are extracted. The temperature after the median filter (magenta line in right upper panel) and the averaged curve (blue line in right lower panel) are also shown. The time course of the YFP/CFP ratio of the AIB neuron (light red line in right middle panel) was calculated from YFP and CFP fluorescences, and the ratio after the median filter is also shown (red line in right middle and lower panel). (C) Plots of the biases of curves (left panel) and the reversal turn frequency (right panel) representing the averages as a function of the entry direction θ. The data obtained on the multi-worm tracker (black lines, n ≥ 12) and on the single-worm tracker (colored lines, n ≥ 14) are shown. (D and E) Plots of the biases of curves (D) and the reversal turn frequency (E) representing the averages as a function of the standardized ratio change (see Materials and methods) of AIB (upper panels). Standardized ratio change was divided into “High activity” or “Low activity” according to the median value (Figure 7G), and the curving bias and the reversal turn frequency were averaged within High (orange columns in lower panels) or Low (cyan columns in lower panels) activity while the animals are moving up or down the thermal gradient. Dashed line in right lower panel shows the average of the reversal turn frequency on the constant temperature at 23°C obtained in MWT. (F) Fractional histogram showing the standardized ratio change of AIB while the animals are moving up the thermal gradient (deep red) and those while moving down the thermal gradient (deep blue). (G) Comparison of the average (left panel), the median (middle panel), and the variance (right panel) of the standardized ratio change of AIB between while the animals are moving up the thermal gradient (deep red columns) and those while moving down the thermal gradient (deep blue columns). n ≥ 14. Error bars indicate SEM. Friedman rank sum test together with repetitive Wilcoxon signed rank tests (D); pairwise test for multiple comparisons using Holm’s method (E); paired Student’s t-test (G). **p < 0.01, *p < 0.05.

## Discussion

In this study, we show that thermotaxis is generated through overlapping but distinct sets of neural circuits, each of which differentially biases individual behavioral components depending on the context of the animals’ position on a temperature gradient relative to the cultivation temperature. These observations can be considered within the framework of degeneracy (Edelman and Gally, 2001; Tononi et al., 1999), which theoretically provides both robustness and flexibility in a system (Wagner, 2005; Whitacre, 2010).

A system with degeneracy is composed of multiple subsystems whose components are partially shared. Consistent with this characteristic, we found that the sub-circuits that differently regulate the behavioral components overlap by sharing several neurons such as AFD, AIY, AIB, AIZ, and SMD (Figures 4, 5, S5-S7). Even after ablating the shared neuron, the regulations of some behavioral components were retained or enhanced, and the cell-ablated animals could still migrate toward the *Tc* (Figure 6). Because of such robustness in the deficient circuits, previous studies might have faced difficulties in identifying and defining neural circuits for thermotaxis (Beverly et al., 2011; Luo et al., 2014a). Similar problems might have appeared in the studies on other types of behavior such as isothermal tracking (Mori and Ohshima, 1995), chemotaxis (Guillermin et al., 2017; Iino and Yoshida, 2009; Luo et al., 2014b), and exploratory behavior (Gray et al., 2005). Also in crustacean and mammalian brain networks, robustness is an obstacle for inferring functional connections that can only be resolved by applying statistical methods (Schwab et al., 2010; Srinivasan and Stevens, 2011). Our observations suggest that by subdividing the behavior into behavioral components (Figure 2B), we can assess the contribution of individual neurons and link the nervous system with the behavior.

In a system that implements degeneracy, individual subsystems have independent contributions from each other to the entire output (Tononi et al., 1999). Therefore, if the subdivisions of behavior are appropriate, the relative sub-circuits might be identified. In our study, the sub-circuits that mediate curves, the speed, and reversal turns are relatively well-defined (Figures 4D, 4H and 5B), suggesting that these subdivisions were successful. By contrast, the sub-circuits mediating shallow turns and omega turns were less defined (Figures 4F and S7B), suggesting that other classifications of turns might be needed to fully elucidate their underlying circuitry as some possibilities have been previously proposed (Broekmans et al., 2016; Kim et al., 2011). Also, non-rule based classifications of behaviors (Brown et al., 2013; Yamaguchi et al., 2018), especially the description of the state of the animal in shape space (Stephens et al., 2008), are reported to be successful in assessing the impact of cell-ablations (Hums et al., 2016; Yan et al., 2017) and might enable the definition of further sub-circuits.

We observed the context-dependent employments of the different sets of the behavioral components (Figures 3C and 3D) under the *T<T_c_* or *T>T_c_* conditions. Given such context-dependent aspects in a system, which is another characteristic of degeneracy, investigation of a system should be better performed under appropriate contexts relevant to a behavior in order to provide mechanistic insights of how a system operates. This might explain several disparities among previous studies on thermotaxis (Hedgecock and Russell, 1975; Luo et al., 2014a; Mori and Ohshima, 1995; Ohnishi et al., 2011; Ryu and Samuel, 2002). Some of these disparities were shown to be due to the difference of the thermal environments (Jurado et al., 2010; Ramot et al., 2008). On steep thermal gradients or in the temperature region distant from the *T_c_*, the animals migrate toward the *T_c_* in the *T>T_c_* condition but not in the *T<T_c_* condition. These features might reflect the context-dependent strategies of the nervous system observed in this study; the curves were mainly employed in the *T<T_c_* condition and the reversal turns were mainly employed in the *T>T_c_* condition (Figure 3). Indeed, the biases of the curves were weakened on the steep thermal gradient (Figure S3) and disappeared in the region distant from the *T_c_* (Figure S4). These observations tell us that the discrimination of contexts is a critical step to investigate the nervous system.

We also observed the context-dependent behavioral regulation by a single interneuron AIB (Figures 7D and 7E) under the dT/dt<0 or dT/dt>0 situations. Considering that the difference of the AIB activity itself was not observed under the two situations (Figures 7F and 7G) and that high (or low) activity of AIB is associated with a bias of the curves (or frequency of the reversal turns) under the dT/dt>0 (or dT/dt<0) but not under the dT/dt<0 (or dT/dt>0) situation, the context-dependent regulations of the curves (or reversal turns) might be achieved by the dynamics of the entire sub-circuits (Figures 4D and 5B). In the neural networks of *Drosophila* larvae, for example, the ratio of neural activities in a circuit is shown to be critical for explaining behavioral choices (Jovanic et al., 2016). In addition, our calcium imaging analyses suggest that the different ranges of the AIB activity regulate the distinct behavioral components. Another first-layer interneuron AIY is indeed known to encode reversals and speeds through digital- and analog-like activities, respectively (Li et al., 2014). These context-dependent behavioral regulations by the different activity patterns of a single neuron might be a prevalent strategy for the nervous system to implement degeneracy.

Our MWT analyses demonstrated that the flexible switching between the migrations up and down the thermal gradient is achieved by the opposed bias of various behavioral components under the *T<T_c_* or *T>T_c_* conditions (Figures 2 and 3). For example, when the animals are moving up the thermal gradient in the *T<T_c_* condition, the turning frequencies were lower, whereas when moving up in the *T>T_c_* condition, the turning frequencies were higher (Figure 2C). One candidate source for this flexibility is the neurotransmission from sensory neurons to interneurons. Several studies have reported that a single sensory neuron can evoke different kind of responses in an identical interneuron through glutamatergic and/or peptidergic transmissions (Guillermin et al., 2017; Kuhara et al., 2011; Narayan et al., 2011; Tsunozaki et al., 2008). Such alterations in synaptic activity can drive opposite behaviors in response to identical stimuli (Cho et al., 2016; Guillermin et al., 2017; Hawk et al., 2018). Another candidate source is the effects of feedback from downstream neurons. In interneuron and motor neuron layers, the motor command sequences are always represented even when the animals are not moving (Hendricks et al., 2012; Kato et al., 2015; Wen et al.,2012). Therefore, the activities of the upstream interneurons could be modulated by those pervasive dynamics. Indeed, the response of AIB to odor stimuli via the AWC sensory neuron is affected by the state of the downstream interneurons RIM and AVA (Gordus et al., 2015). Some studies have suggested that sensory inputs could be converted into appropriate motor outputs after being integrated with those dynamics (Hendricks and Zhang, 2013; H. Liu et al., 2018). Also in mammalian brains, feedback from downstream neurons are known to play an important role in the visual system (Pascual-Leone and Walsh, 2001) or somatosensory system (Manita et al., 2015). These interactions with pervasive dynamics might result in the context-dependent recruitments of the different sub-circuits (Figures 4D, 4E and 4H). Our analysis indicated that the different class of head motor neurons, SMD and RME, were recruited differently to regulate the curves and the shallow turns under the *T<T_c_* or *T>T_c_* conditions. Since these neurons are excitatory and inhibitory neurons, respectively, such context-dependent recruitments of different head motor neurons might enable the opposed regulations of the behavioral components.

Our study shows that the implementation of circuit degeneracy may be a prevalent strategy for the nervous system to execute robust and flexible behavior, which is a sophisticated aspect of animal behavior.

## Materials and Methods

### Strains

*C. elegans* animals were cultivated under the standard condition (Brenner, 1974). Adult hermaphrodites were used in this study. N2 (Bristol) was the wild-type strain, and all the other strains used in this study were derived from N2.

Cell-ablated strains were generated by the expression of reconstituted caspases (Chelur and Chalfie, 2007) and by mito-miniSOG with the FLP/FRT strategy (Davis et al., 2008; Qi et al., 2012). Each of the plasmids expressing reconstituted caspase was injected at 25 ng/μl and each of the plasmids expressing mito-miniSOG was injected at 50-75 ng/μl, and in both cases, pKDK66 (ges-1p::NLS::GFP) (50 ng/μl) was co-injected as an injection marker. Cell-specific expressions were achieved by using the promoter sets listed in Table S1. Specificity was confirmed by expressing TagRFP under the listed promoter sets with the FLP/FRT strategy and also by checking the fluorescence from miniSOG. Extrachromosomal arrays were integrated into the genome via gamma irradiation-induced mutagenesis except for *njIs127*, which was spontaneously generated through the daily maintenance, and outcrossed more than four times before analysis. PY7505 was kindly provided by Piali Sengupta, Brandeis University, MA, USA (Beverly et al., 2011).

Integrants of recCaspasese were crossed into integrated reporter lines listed in Table S2 that express GFPs or TagRFPs in several neurons including the neuron of interest. Losses of neurons were confirmed at the adult stage by the disappearance of fluorescence from the reporter proteins. Plates containing OP50 and the L1 stage animals expressing mito-miniSOG were exposed, without any covers, to pulsed blue light (488 nm) in 0.5 sec on and 1.5 sec off cycles for 30 min. The blue light intensity received by the animals was measured as 106 mW/cm^2^. Losses of neurons were confirmed at the adult stage by the disappearance of fluorescence from the miniSOG. To control the shutters, we used five spot-type deep UV lamps (SP-9250EF-N, USHIO) connected by light guide fiber units (SP-155XQ-S11, USHIO) and a control box (SP-SC-N, USHIO). The efficiencies of the cell ablations are also listed in Table S1.

### Thermotaxis Assay

Thermotaxis (TTX) assays were performed as previously described (Ito et al., 2006). Animals cultivated at 17 °C, 20 °C, or 23 °C were placed on the center of the assay plate (13.6 cm×9.6 cm, 1.45 cm height) containing 18 ml of TTX medium with 2% agar, and were allowed to freely move for 60 min. The center of the plate was adjusted at 14 °C, 17 °C, 20 °C, 23 °C, and 26 °C depending on the experiments. The plate was maintained with a linear thermal gradient of approximately 0.45 °C/cm.

### Behavioral Recording

Behavioral recordings were performed using a Multi-Worm Tracker (Swierczek et al., 2011; Yamaguchi et al., 2018) with a CMOS sensor Camera Link Camera (8 bits, 4,096 × 3,072 pixels; CSC12M25BMP19-01B, Toshiba-Teli), a lens adaptor (F-TAR2), a Line-Scan Lens (35mm, f/2.8; YF3528, PENTAX), and a PCIe-1433 camera-link frame grabber (781169-01, National Instruments). The camera was mounted at a distance above the assay plate resulting in an image with 33.2 μm per pixel. The frame rate of recordings was approx. 13.5 Hz. Images were captured and processed by custom software written in LabView (National Instruments) and a custom image analysis library written in C++, which detect animals and measure parameters such as the positions and the postures of animals.

### Behavioral Analysis

The MWT system automatically identifies animals and provides the positions of their centers of mass and the 11 points along their bodies, as well as their body lengths, widths, and so on (Swierczek et al., 2011). Using these data, the behavioral analysis was performed with a custom-built MATLAB (MathWorks) script. For each frame, we defined the moving direction as the vectors from the current centroid to the following centroid (1 sec after), and calculated the *curve* by the angle between the previous moving direction (1 sec before) and the current moving direction. When an animal performs the *omega turn*, its head and tail become close together accompanying the decrease of the estimated body length in the system. Therefore, if the body length was estimated shorter than 1.5 standard deviation from the mean and the curve value at that time was greater than 90°/sec, we regarded the animal as performing the omega turn. To detect *shallow turns*, we defined the head swing for each frame as the angle between the vector from the 3rd point to the 1st point and the vector from the 7th point to the 5th point along the worm’s body. If the head swing was over 2 standard deviation from the mean and the curve value at that time was in the range of 15-90°, we regarded the worm as performing the shallow turn. *Reversals* were detected by the smoothed curve (the moving average of the curves within three frames) which was greater than 150°/sec. If a reversal was followed by an omega turn within 6 seconds, these two components were combined into a *reversal turn* (Iino and Yoshida, 2009; Pierce-Shimomura et al., 1999). All the curve thresholds described above were determined following the previous proposals (Kim et al., 2011; Schild and Glauser, 2013).

### Computer Simulation

Thermotaxis behavior was simulated using another custom-built MATLAB script. For each simulation, 100 animals were run sequentially. Animals were considered as dimensionless points in a 13.6 cm (x axis) × 9.6 cm (y axis) plate, with a linear thermal gradient from 14 to 20 °C for the *T<Tc* condition and from 20 to 26 °C for the *T>Tc* condition along the x axis. Animals started from the center of a plate, while y coordinates and initial directions were randomized. For every second, animals decided whether to do an omega turn, a shallow turn, a reversal, a reversal turn, or a curve (Figure 3A). Event probabilities of each behavioral component were defined according to the experimental data of turning frequencies. When animals decided to do any turns, the next moving directions (θ) were defined according to the experimental data of exit directions (Φ). The next positions (x, y) were defined together with the experimental data of the displacements during the individual turns that were also calculated in MWT analysis. When animals decided to do a curve, the next moving directions θ were defined according to the experimental data of curving biases (φ). The next positions (x, y) were defined together with the experimental data of the speed. If an animal reaches the plate border, it was set to do specular reflection. When disabling each of the behavioral components (Figure 3C), we replaced the experimental data of interest with the data taken from the animals on the constant temperature. When enabling each of the behavioral components, we replaced the experimental data of interest on the constant temperature with the data taken from the animals on the thermal gradient. When exchanging each of the behavioral components (Figure 6G), we replaced the experimental data of interest with the data taken from the wild-type animals on the thermal gradient, while the probability of a curve (Figure 3A) was kept same. Every experimental data was applied as a function of moving direction θ. Besides, different data set were applied depending on whether the animals were on the fraction 1 −2, the fraction 3–6, or the fraction 7–8 of a thermotaxis plate (Figure 1C). Each simulation lasts for 30 min, and the simulations were iterated 100 times and the time courses of TTX indices were averaged within them.

### Calcium Imaging in Freely Moving Animals

Ca^2+^ imaging recordings in freely moving animals were performed as previously described (Tsukada et al., 2016) with custom modifications. The FRET-based calcium probe yellow cameleon X 1.60 was expressed in the neuron of interest (Table S2). A dual-view was equipped with 05-EM CFP/YFP (505 dcxr) filter cube (Molecular Devices), and images were acquired using EM-CCD camera (C9100-13, Hamamatsu Photonics). Simultaneous tracking was performed using a CMOS camera (Grasshopper Express GX-FW-28S5M-C, FLIR Integrated Imaging Solutions) at 30 frames per second with continuous halogen illumination (TH4-100, Olympus) through an IR filter (IR-76, Fujifilm). Along the trajectory of animals (Figures 7B and S8B), behavioral components were detected in a similar way to that in the MWT analysis, and the same regulations of the curves and the reversal turns were observed (Figures S7B and S7C).

### Imaging Analysis

The image processing program for the tracking data was written in MATLAB. A neuronal region was defined according to the peak intensity and size (9 pixels) in an YFP image. In each image, the averaged background intensity within 9 pixels was subtracted from the average fluorescence intensities of the neuronal regions. Intercellular calcium concentration change was estimated by taking the YFP/CFP fluorescence ratio *(Ratio)* and YFP/CFP ratio change *(Ratio change)*, which was normalized within each assay *(Standardized ratio change*) to compare the activity among the assays. A median filter within a moving 15 sec temporal window was applied to the time course ratio to eliminate the noise independent from calcium signal. The *averaged curve* was calculated by averaging the curves within a moving 90 sec temporal window. For the analysis of the relationship between the neural activities and the regulations of the behavioral components, we defined an activity that was higher than the median of all the activities within an assay as a *High activity* and an activity that was lower than the median as a *Low activity*.

### Quantification and Statistical Analysis

Experimental data are expressed as mean ± SEM. Simulation data are expressed as mean. For comparison of the data from behavioral analysis in MWT, we used a paired Student’s í-test and a one-way ANOVA followed by a Tukey–Kramer post hoc multiple comparisons test. For comparison of the data from imaging experiments, we used a paired Student’s í-test, pairwise test for multiple comparisons using Holm’s method, and Friedman rank sum test together with repetitive Wilcoxon signed rank tests as noted in the figure legends. Bartlett’s test was used to check for differences in variance among the groups. A difference is considered significant at a value of **p < 0.01 or *p < 0.05.

## Acknowledgements

We thank Martin Chalfie for the reconstituted caspases construct (Addgene plasmids #16082 and #16083); Yishi Jin for the mito-miniSOG construct; Seika Takayanagi-Kiya for the advice on the usage of mito-miniSOG; Eduardo Izquierdo for the advice on the assessment of the simulation; Erik Jorgensen for the FLP and FRT construct; Piali Sengupta for the ASI-ablated strain; Shin Takagi for the unpublished AVE marker strain; Kaveh Ashrafi for the *mgl-1* promoter; Mario de Bono for the *npr-4a* promoter; and Shawn Xu for the information about the *sío-3* promoter. M.I. was supported by KAKENHI 16J05770, A.C.G. by JSPS Postdoctoral Fellowship PE12065, and this work was supported by KAKENHI 16H02516 (to I.M.).

---

**Figure S1.**
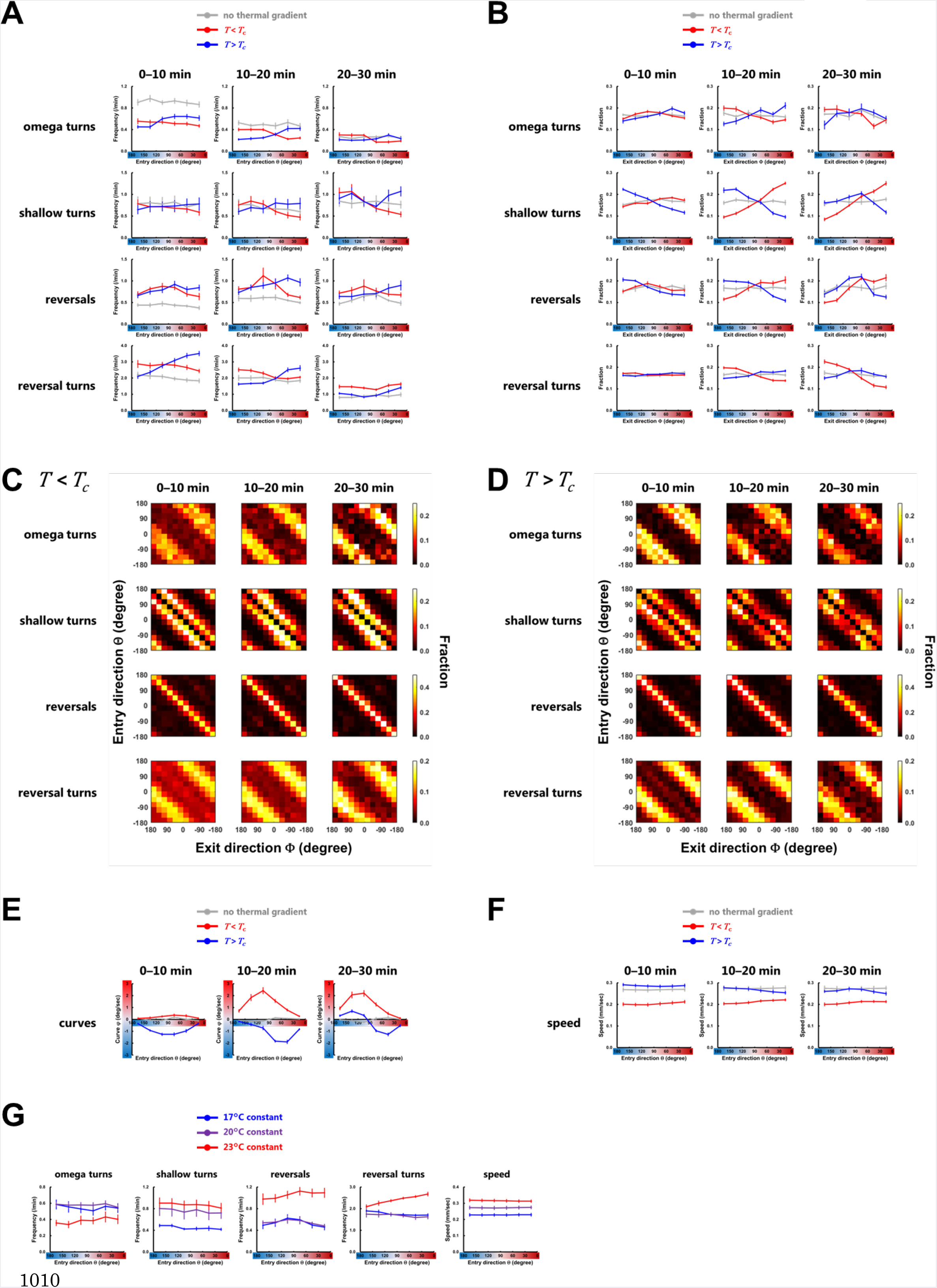
Time course of the regulations of the behavioral components in wild-type animals. (A, B, E, F) Regulations of the behavioral components on the constant temperature (gray lines), in the *T<T_c_* condition (red lines), and in the *T>T_c_* condition (blue lines). (A) Frequency plots of the turns and the reversals representing the averages as a function of the entry direction θ. (B) Fraction plots of the exit direction Φ after the turns and the reversals. (E) Plots of the biases φ of curves representing the averages as a function of the entry direction θ. (F) Speed plots representing the averages as a function of the entry direction θ. (C and D) Heat map of fractions of the exit direction Φ as functions of the entry direction θ in the *T<T_c_* condition (C) and in the *T>T_c_* condition (D). Both θ and Φ are signed to distinguish whether the exit angle is directed toward the upper half or the lower half of assay plates. (G) Frequency plots of the turns and the reversals and speed plots representing the averages as a function of the entry direction θ on the constant temperature at 17°C (blue lines), 20°C (purple lines), and 23°C (red lines). In (A-G), the averages over 0-10 min (left columns), over 10-20 min (middle columns), and over 20-30 min (right columns) are shown.

**Figure S2.**
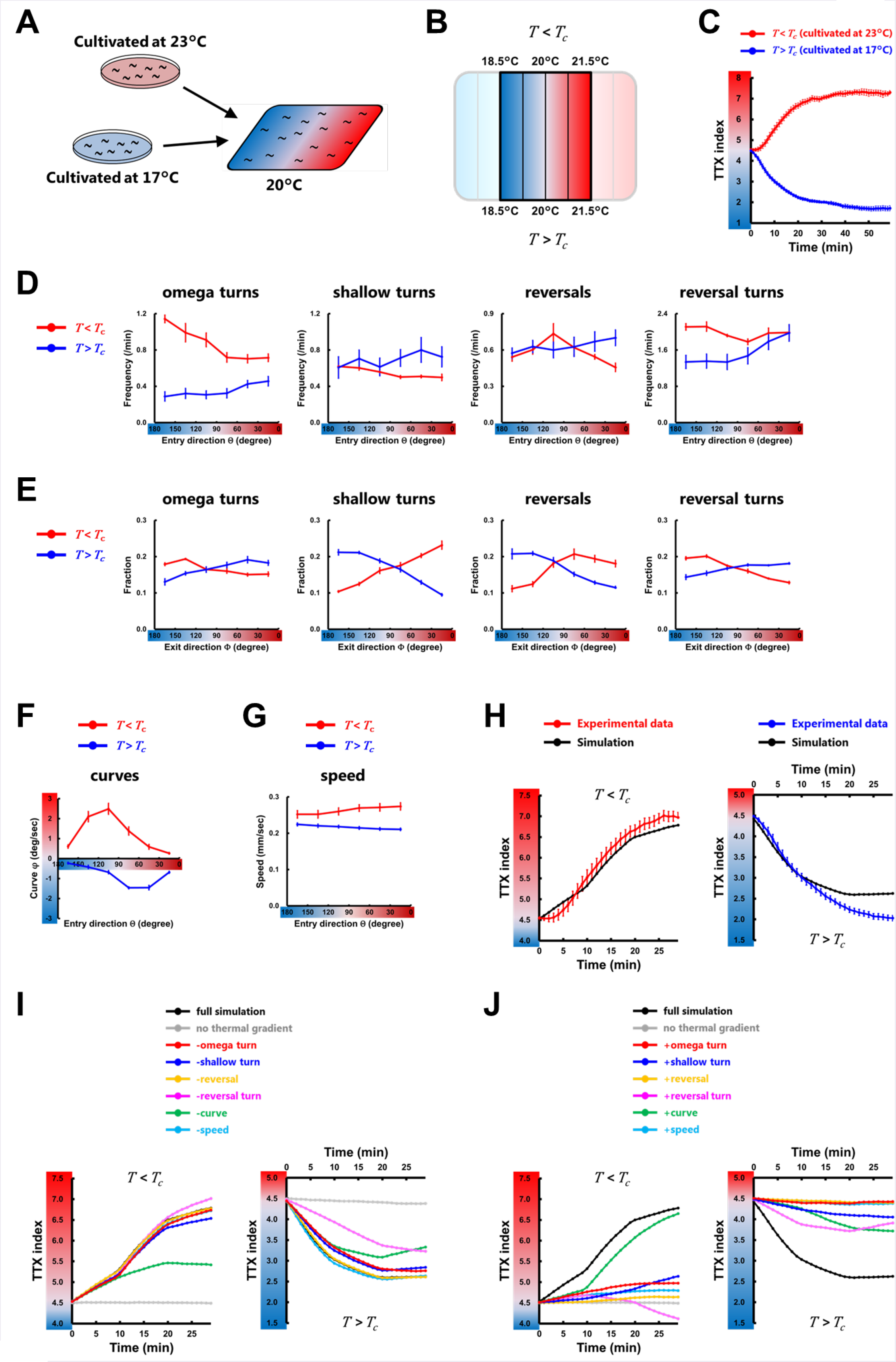
Behavioral components are employed differently depending on the *T_c_*. (A) Animals cultivated at 17°C or 23°C were placed on a thermal gradient with 20°C at the center. The plate was maintained with a linear thermal gradient of approximately 0.45 °C/cm. (B) Temperature range within which the behavioral components were analyzed. (C) The time course of TTX indices in the *T<T_c_* condition (red line) and in the *T>Tc* condition (blue line). (n = 6). (D-G) Regulations of the behavioral components in the *T<T_c_* condition (red lines) and in the *T>T_c_* condition (blue lines). (D) Frequency plots of the turns and the reversals representing the averages as a function of the entry direction θ. (E) Fraction plots of the exit direction Φ after the turns and the reversals. (F) Plots of the biases φ of curves representing the averages as a function of the entry direction θ. (G) Speed plots representing the averages as a function of the entry direction θ. (H) The time course of TTX indices in the simulations (black lines) and that obtained from experimental data (colored lines). (I and J) black lines correspond to the simulation in which all the data of wild-type animals determined by the experiment with the thermal gradient were used, and gray lines correspond to the simulation in which all the data of wild-type animals without the gradient were used. The other colored lines correspond to the simulation with the replacements of the individual behavioral component: the omega turn (red lines), the shallow turn (blue lines), the reversal (yellow lines), the reversal turn (magenta lines), the curve (green lines), and the speed (light blue lines).

**Figure S3.**
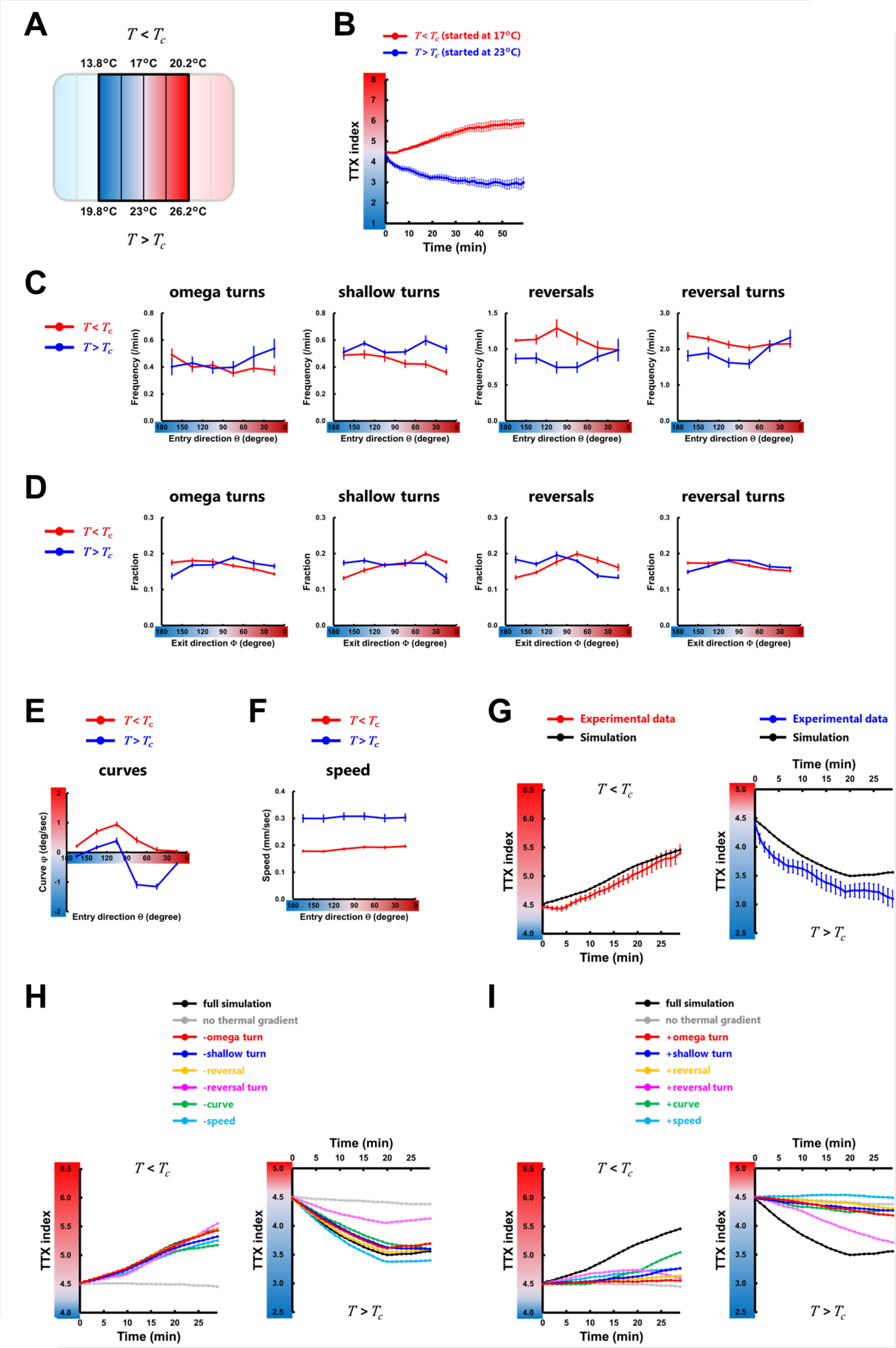
Employments of the behavioral components are slightly affected by the steepness of thermal gradient. (A) Temperature range within which the behavioral components were analyzed. The plate was maintained with a linear thermal gradient of approximately 0.95 °C/cm. (B) The time course of TTX indices in the *T<T_c_* condition (red line) and in the *T>T_c_* condition (blue line). (n ≥ 6). (C-F) Regulations of the behavioral components in the *T<T_c_* condition (red lines) and in the *T>T_c_* condition (blue lines). (C) Frequency plots of the turns and the reversals representing the averages as a function of the entry direction θ. (D) Fraction plots of the exit direction Φ after the turns and the reversals. (E) Plots of the biases φ of curves representing the averages as a function of the entry direction θ. (F) Speed plots representing the averages as a function of the entry direction θ. (G) The time course of TTX indices in the simulations (black lines) and that obtained from experimental data (colored lines). (H and I) black lines correspond to the simulation in which all the data of wild-type animals determined by the experiment with the thermal gradient were used, and gray lines correspond to the simulation in which all the data of wild-type animals without the gradient were used. The other colored lines correspond to the simulation with the replacements of the individual behavioral component: the omega turn (red lines), the shallow turn (blue lines), the reversal (yellow lines), the reversal turn (magenta lines), the curve (green lines), and the speed (light blue lines).

**Figure S4.**
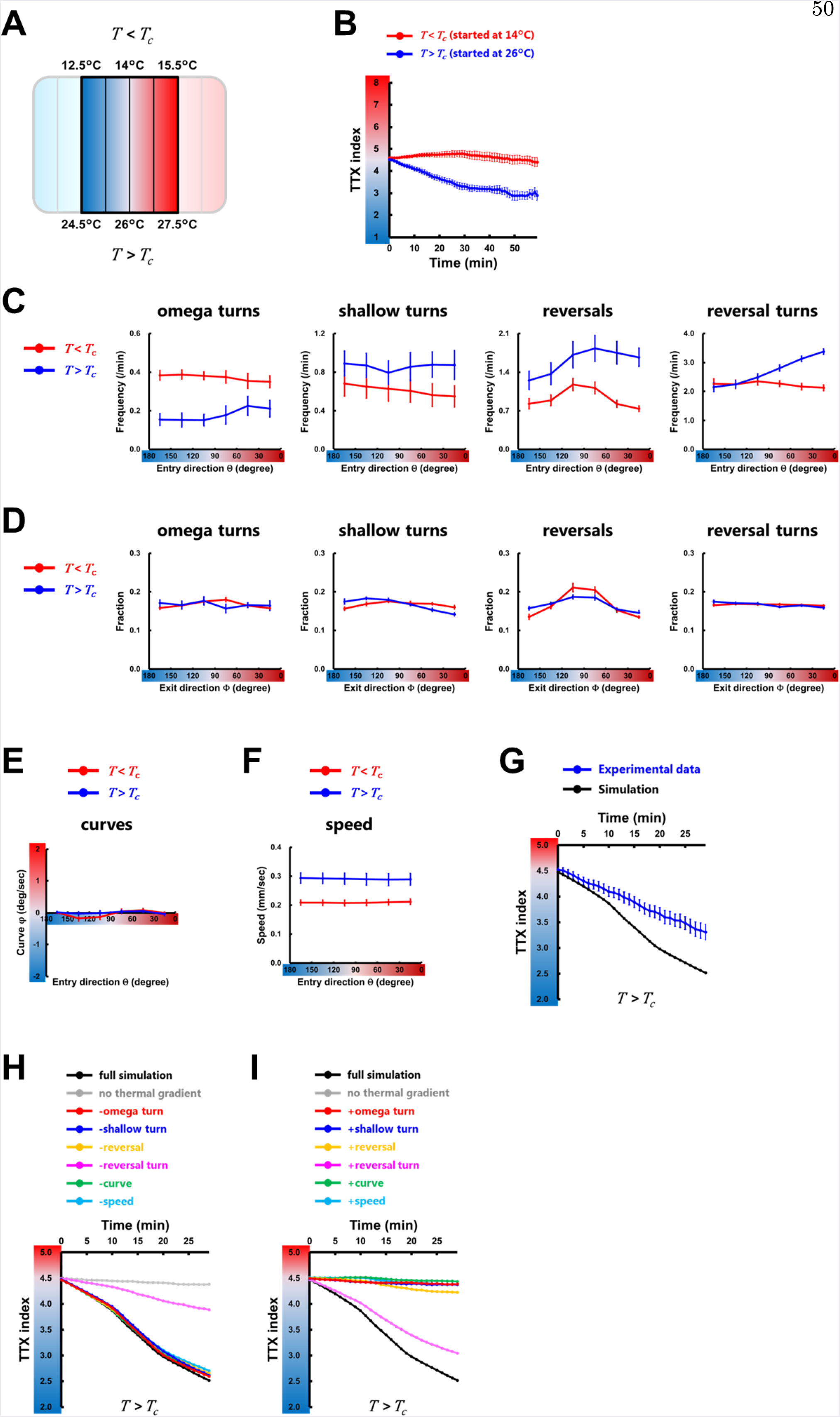
Behavioral components except for the reversal turns are not employed in the region distant from the *T_c_*. (A) Temperature range within which the behavioral components were analyzed. The plate was maintained with a linear thermal gradient of approximately 0.45 °C/cm. (B) The time course of TTX indices in the *T<T_c_* condition (red line) and in the *T>T_c_* condition (blue line). (n ≥ 6). (C-F) Regulations of the behavioral components in the *T<T_c_* condition (red lines) and in the *T>T_c_* condition (blue lines). (C) Frequency plots of the turns and the reversals representing the averages as a function of the entry direction θ. (D) Fraction plots of the exit direction Φ after the turns and the reversals. (E) Plots of the biases φ of curves representing the averages as a function of the entry direction θ. (F) Speed plots representing the averages as a function of the entry direction θ. (G) The time course of TTX indices in the simulations (black line) and that obtained from experimental data (blue line). (H and I) black lines correspond to the simulation in which all the data of wild-type animals determined by the experiment with the thermal gradient were used, and gray lines correspond to the simulation in which all the data of wild-type animals without the gradient were used. The other colored lines correspond to the simulation with the replacements of the individual behavioral component: the omega turn (red lines), the shallow turn (blue lines), the reversal (yellow lines), the reversal turn (magenta lines), the curve (green lines), and the speed (light blue lines).

**Figure S5.**
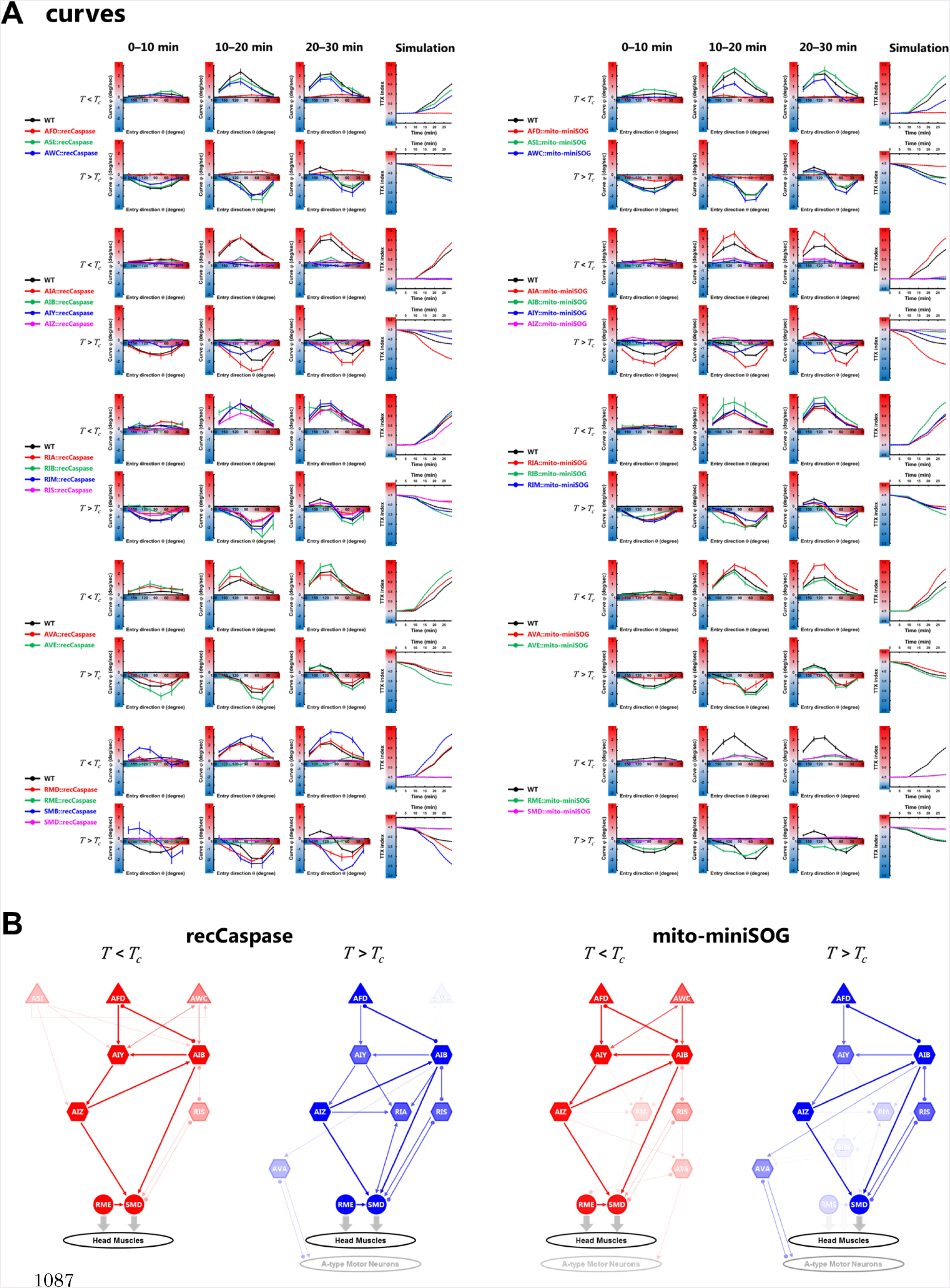
Time course of the regulations of the behavioral components in cell-ablated animals. (A) Plots of the biases φ of curves representing the averages as a function of the entry direction θ and the time course of TTX indices in the simulations. (B) Predicted neural circuits for regulating the curves. (C) Fraction plots of the exit direction Φ after the shallow turns and the time course of TTX indices in the simulations. (D) Predicted neural circuits for regulating the shallow turns. (E) Speed plots and the time course of TTX indices in the simulations. (F) Predicted neural circuits for regulating the speeds. In (A, C, E), the averages over 0-10 min (first columns), over 10–20 min (second columns), over 20–30 min (third columns), and the time course of TTX indices in the simulations (fourth columns) are shown. n ≥ 5. In (B, D, F), predicted neural circuits in the *T<T_c_* condition (red) and in the *T>T_c_* condition (blue) are shown. The thickness and color strength of each neuron represent the functional importance of the neuron predicted from the analysis and were determined as follows: For each neuron, the differences between the migration index of the wild-type animals and the index of the cell-ablated animals expressing reconstituted caspases (left panels) or mito-miniSOG (right panels) were calculated. The difference is used to determine the color strength, where the color strength of each neuron is proportional to this value. The color strength of each line is identical to the strength of the color of one of the two connected neurons with lower strength, and the thickness of each line is proportional to this color strength. In RIS, SMB, and RMD neurons, we applied the data of the animals expressing recCaspase on both panels because we could not obtain the animals expressing mito-miniSOG specifically in these neurons.

**Figure S6.**
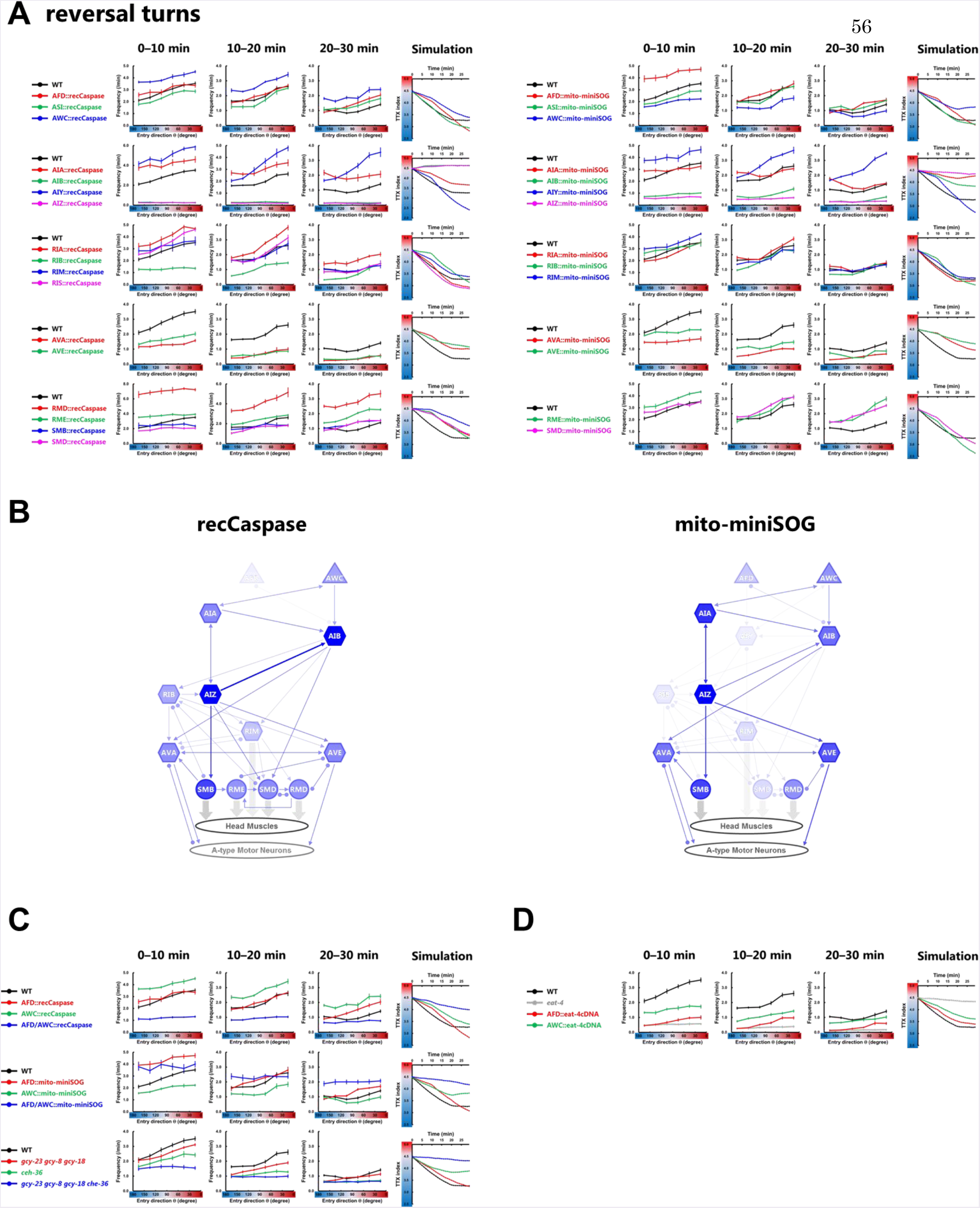
The time course of the regulations of the reversal turns in cell-ablated animals and cell-deficient mutants. (A, C, D) Plots of the reversal turn frequency representing the averages as a function of the entry direction θ and the time course of TTX indices in the simulations. (A) Cell-ablated animals. (C) AFD-AWC double ablated/deficient animals. (D) Glutamate transporter-deficient mutant *eat-4*, the mutant expressing the wild-type form of an *eat-4* cDNA only in AFD, and the mutant expressing an *eat-4* cDNA only in AWC. The averages over 0-10 min (first columns), over 10-20 min (second columns), over 20-30 min (third columns), and the time course of TTX indices in the simulations (fourth columns) are shown. n ≥ 5. (B) Predicted neural circuits for regulating the reversal turns. In RIS, SMB, and RMD neurons, we applied the data of the animals expressing recCaspase on both panels because we could not obtain the animals expressing mito-miniSOG specifically in these neurons.

**Figure S7.**
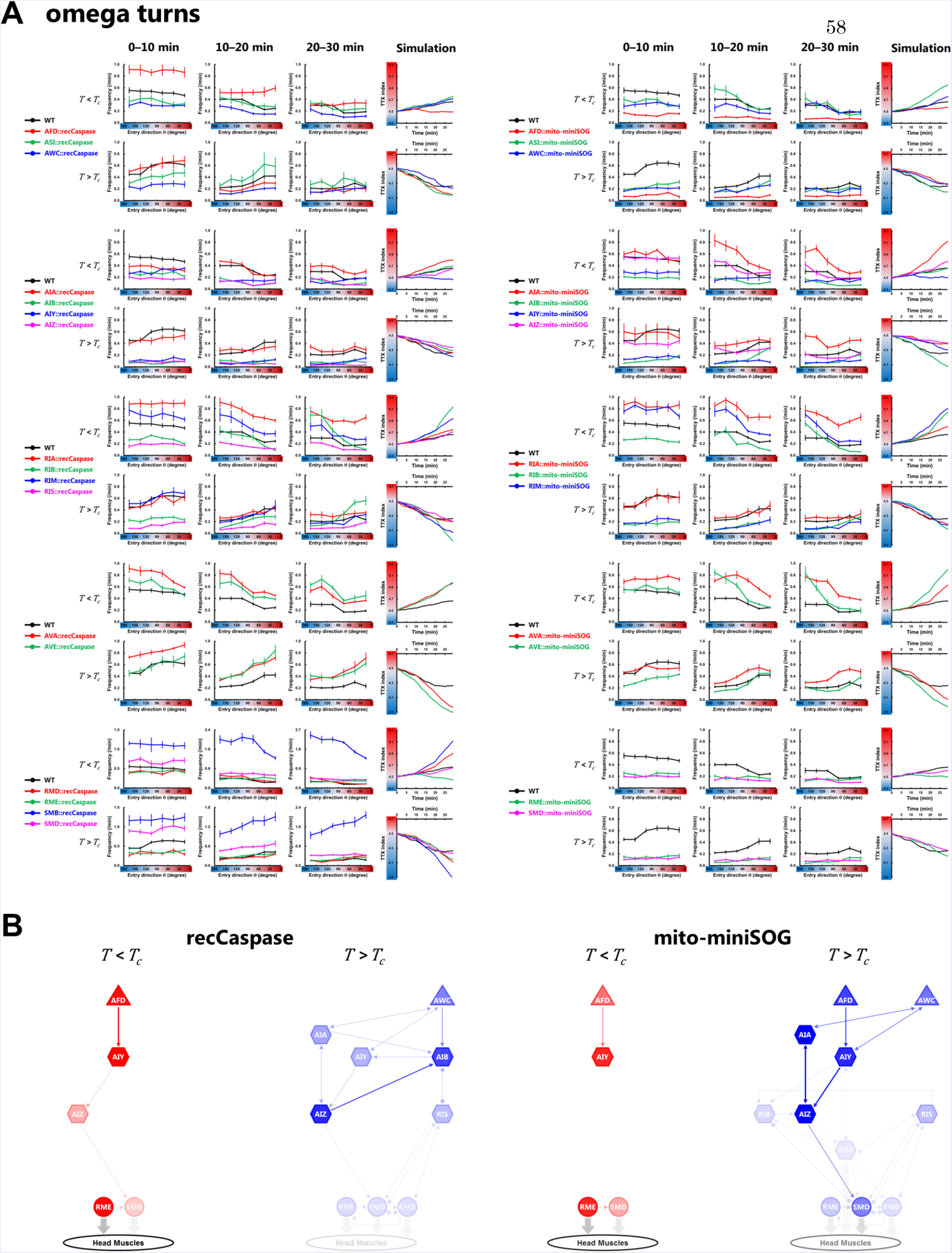
The time course of the regulations of the omega turns in cell-ablated animals and cell-deficient mutants. (A) Plots of the omega turn frequency representing the averages as a function of the entry direction θ and the time course of TTX indices in the simulations. The averages over 0-10 min (first columns), over 10-20 min (second columns), over 20–30 min (third column), and the time course of TTX indices in the simulations (fourth columns) are shown. n ≥ 5. (B) Predicted neural circuits for regulating the omega turns in the *T<T_c_* condition (red) and in the *T>T_c_* condition (blue). In RIS, SMB, and RMD neurons, we applied the data of the animals expressing recCaspase on both panels because we could not obtain the animals expressing mito-miniSOG specifically in these neurons.

**Figure S8.**
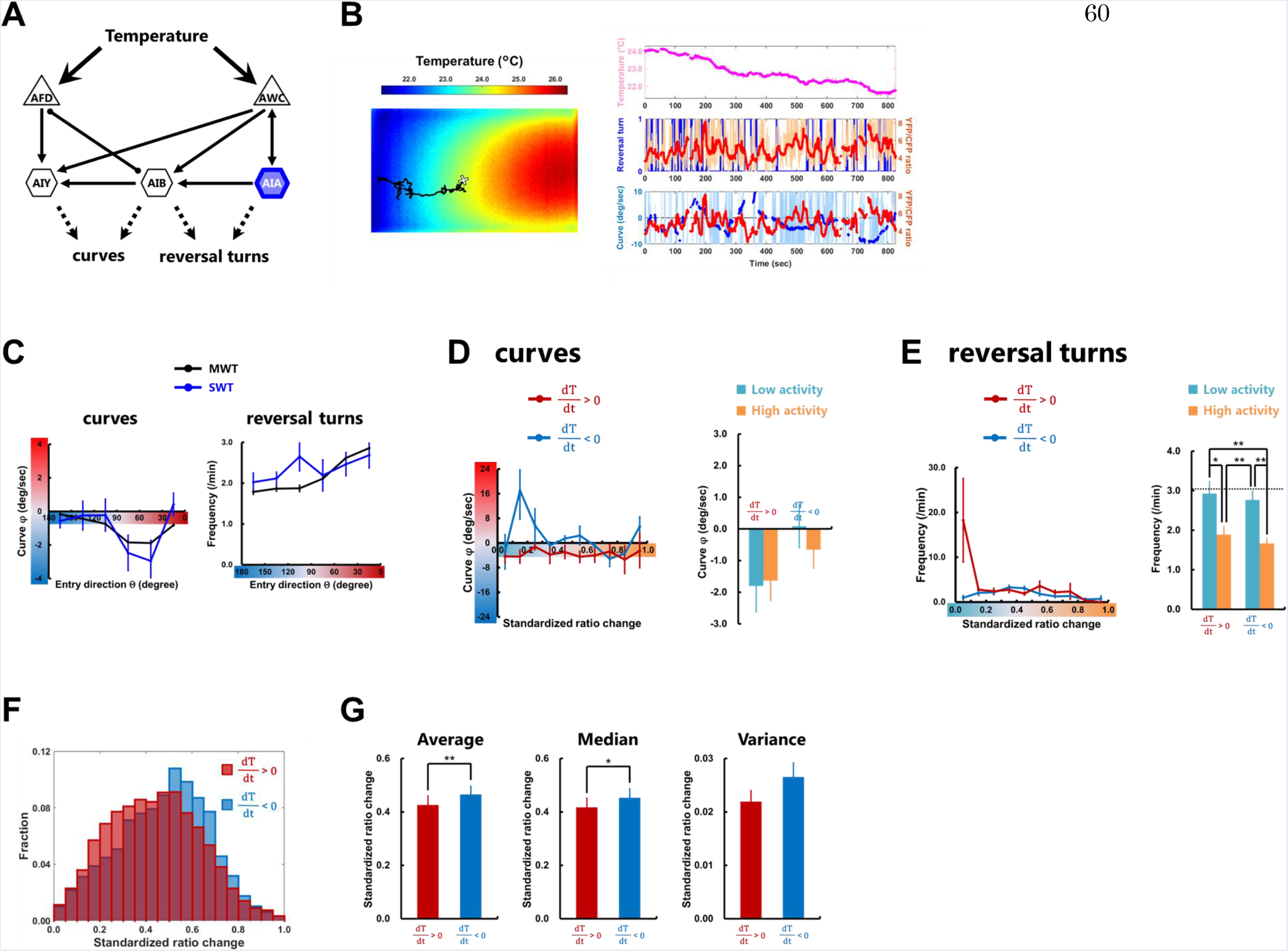
AIA neuron regulates the reversal turns regardless of context. (A) Representative sensory neurons and first-layer interneurons, which regulate the reversal turns or the curves. Thin arrows indicate chemical synapses and an undirected line with round endings indicates gap junction. (B) Representative thermography image (left panel) taken together with calibrating temperature measurements using a thermocouple sensor. Projection of a trajectory (black line) shows how the animal searches the thermal environment. The white + marks the starting point when recording starts. From this trajectory, the time course of the temperature changes is obtained (light magenta line in right upper panel), and the reversal turn (blue line in right middle panel) and the curve (light blue line in right lower panel) are extracted. The temperature after the median filter (magenta line in right upper panel) and the averaged curve (blue line in right lower panel) are also shown. The time course of the YFP/CFP ratio of the AIA neuron (light red line in right middle panel) was calculated from YFP and CFP fluorescences, and the ratio after the median filter is also shown (red line in right middle and lower panel). (C) Plots of the biases of curves (left panel) and the reversal turn frequency (right panel) representing the averages as a function of the entry direction θ. The data obtained on the multi-worm tracker (black lines, n ≥ 12) and on the single-worm tracker (colored lines, n ≥ 14) are shown. (D and E) Plots of the biases of curves (D) and the reversal turn frequency (E) representing the averages as a function of the standardized ratio change (see Materials and methods) of AIA (upper panels). Standardized ratio change was divided into “High activity” or “Low activity” according to the median value (Figure S8G), and the curving bias and the reversal turn frequency were averaged within High (orange columns in lower panels) or Low (cyan columns in lower panels) activity while the animals are moving up or down the thermal gradient. Dashed line in right lower panel shows the average of the reversal turn frequency on the constant temperature at 23°C obtained in MWT. (F) Fractional histogram showing the standardized ratio change of AIA while the animals are moving up the thermal gradient (deep red) and those while moving down the thermal gradient (deep blue). (G) Comparison of the average (left panel), the median (middle panel), and the variance (right panel) of the standardized ratio change of AIA between while the animals are moving up the thermal gradient (deep red columns) and those while moving down the thermal gradient (deep blue columns). n ≥ 15. Error bars indicate SEM. Pairwise test for multiple comparisons using Holm’s method (D); Friedman rank sum test together with repetitive Wilcoxon signed rank tests (E); paired Student’s t-test (G). **p < 0.01, *p < 0.05.

**Table S1.** Cell-ablated strains used in this study. Efficiencies of cell-ablations by recCaspasese were estimated by crossing the listed lines into integrated reporter lines listed in Table S2 and checking the disappearance of fluorescence from the reporter proteins. *Efficiency of *njIs93* was estimated in the heterozygous state. Efficiencies of cell-ablations by mito-miniSOG were estimaterd by checking the disappearance of fluorescence from the miniSOG after the illumination of blue light at the L1 stage.

| Strain name | Genotype | Experiment | Ablation Efficiency (%) |
| --- | --- | --- | --- |
| IK2809 | *njls80[gcy-8p::cz::caspase-3(p17), gcy-8p::caspase-3(p12)::nz, ges-1p::nls-GFP] (X)* | AFD ablation | 100 |
| IK3048 | *njls89[gcy-8p::tomm-20(N'-55AA)::miniSOG, ges-1p::nls-GFP] (III)* |  | 92.7 |
| PY7505 | *oyls84[gcy-27p::cz::caspase-3(p17), gpa-4p::caspase-3(p12)::nz, gcy-27p::GFP, unc-122p::dsRed]* | ASI ablation | (Beverly et al., 2011) |
| IK3176 | *njls104[gcy-27p::flp, gpa-4p::frr::tomm-20(N'-55AA)::miniSOG, ges-1p::nls-GFP] (IV)* |  | 91.9 |
| IK2808 | *njls79[ceh-36p::cz::caspase-3(p17), ceh-36p::caspase-3(p12)::nz, ges-1p::nls-GFP] (X)* | AWC ablation | 98.4 |
| IK3125 | *njls98[ceh-36p::tomm-20(N'-55AA)::miniSOG, ges-1p::nls-GFP] (I)* |  | 100 |
| IK3263 | *njls120[ins-1p::cz::caspase-3(p17), gcy-28dp::caspase-3(p12)::nz, ges-1p::nls-GFP] (V)* | AIA ablation | 64.6 |
| IK3240 | *njls115[ins-1p::flp, gcy-28dp::frr::tomm-20(N'-55AA)::miniSOG, ges-1p::nls-GFP] (IV)* |  | 74.1 |
| IK3066 | *njls92[inx-1p::cz::caspase-3(p17), odr-2(2b)p::caspase-3(p12)::nz, ges-1p::nls-GFP] (II)* | AIB ablation | 100 |
| IK3388 | *njls131[odr-2(2b)p::flp, inx-1p::frr::tomm-20(N'-55AA)::miniSOG, ges-1p::nls-GFP] (X)* |  | 100 |
| IK2710 | *njls62[AIYp::cz::caspase-3(p17), AIYp::caspase-3(p12)::nz, ges-1p::nls-GFP] (V)* | AIY ablation | 100 |
| IK2962 | *njls87[AIYp::tomm-20(N'-55AA)::miniSOG, ges-1p::nls-GFP] (IV)* |  | 100 |
| IK3179 | *njls107[acc-2p::cz::caspase-3(p17), odr-2(2b)p::caspase-3(p12)::nz, ges-1p::nls-GFP] (IV)* | AIZ ablation | 100 |
| IK3241 | *njls116[acc-2p::flp, odr-2(2b)p::frr::tomm-20(N'-55AA)::miniSOG, ges-1p::nls-GFP] (IV)* |  | 99.7 |
| IK2910 | *njls84[glr-3p::cz::caspase-3(p17), glr-3p::caspase-3(p12)::nz, ges-1p::nls-GFP] (III)* | RIA ablation | 100 |
| IK3289 | *njls123[glr-3p::tomm-20(N'-55AA)::miniSOG, ges-1p::nls-GFP] (V)* |  | 100 |
| IK3049 | *njls90[trp-1p::cz::caspase-3(p17), sto-3p::caspase-3(p12)::nz, ges-1p::nls-GFP] (I)* | RIB ablation | 96.6 |
| IK3238 | *njls113[ser-4p::flp, sto-3p::frr::tomm-20(N'-55AA)::miniSOG, ges-1p::nls-GFP] (II)* |  | 95.5 |
| IK3067 | *njls93[glr-1p::cz::caspase-3(p17), tdc-1p::caspase-3(p12)::nz, ges-1p::nls-GFP] (X)* | RIM ablation | 66.7* |
| IK3144 | *njls100[glr-1p::flp, tdc-1p::frr::tomm-20(N'-55AA)::miniSOG, ges-1p::nls-GFP] (III)* |  | 92.9 |
| IK3325 | *njls126[nlr-1p::cz::caspase-3(p17), ggr-2p::caspase-3(p12)::nz, ges-1p::nls-GFP] (V)* | RIS ablation | 98.5 |
| IK3175 | *njls103[nmr-1p::cz::caspase-3(p17), flp-18p::caspase-3(p12)::nz, ges-1p::nls-GFP] (II)* | AVA ablation | 100 |
| IK3327 | *njls128[flp-18p::flp, nmr-1p::frr::tomm-20(N'-55AA)::miniSOG, ges-1p::nls-GFP] (II)* |  | 82.7 |
| IK3177 | *njls105[nmr-1p::cz::caspase-3(p17), opt-3p::caspase-3(p12)::nz, ges-1p::nls-GFP] (X)* | AVE ablation | 97.8 |
| IK3324 | *njls125[opt-3p::flp, nmr-1p::frr::tomm-20(N'-55AA)::miniSOG, ges-1p::nls-GFP] (X)* |  | 100 |
| IK3178 | *njls106[odr-2(18)p::cz::caspase-3(p17), nep-2p::caspase-3(p12)::nz, ges-1p::nls-GFP] (III)* | SMBD/V ablation | 53.3 |
| IK3326 | *njls127[flp-22p::cz::caspase-3(p17), lgc-55p::caspase-3(p12)::nz, ges-1p::nls-GFP] (V)* | SMDD/V ablation | 75.2 |
| IK3376 | *njls129[lgc-55p::flp, flp-22p::frr::tomm-20(N'-55AA)::miniSOG, ges-1p::nls-GFP] (I)* |  | 95.9 |
| IK3237 | *njls121[glr-1p::cz::caspase-3(p17), mgl-1p::caspase-3(p12)::nz, ges-1p::nls-GFP] (II)* | RMD/D/V ablation | 34.8 |
| IK3404 | *njls132[ser-2(2)p::cz::caspase-3(p17), ntr-2p::caspase-3(p12)::nz, ges-1p::nls-GFP] (X)* | RMED/V ablation | 56.9 |
| IK3377 | *njls130[ser-2(2)p::flp, ntr-2p::frr::tomm-20(N'-55AA)::miniSOG, ges-1p::nls-GFP] (X)* |  | 95.5 |
| IK3242 | *njls117[gcy-8p::cz::caspase-3(p17), gcy-8p::caspase-3(p12)::nz, ges-1p::nls-GFP] (I); njls79* | AFD/AWC ablation | - |
| IK3227 | *njls98; njls89* |  | - |

**Table S2.** Strains carrying cell markers, mutations, or calcium indicators used in this study.

| Strain name | Genotype | Experiment |
| --- | --- | --- |
| IK0673 | *njls2[nhr-38p::GFP, AIYp::GFP] (V)* | AFD/AIY marker |
| IK2952 | *njls86[sra-6p::GFP] (X)* | ASI marker |
| IK2811 | *njls82[ceh-36p::GFP, glr-3p::GFP] (I)* | AWC/RIA marker |
| IK3237 | *njls112[gcy-28dp::GFP] (X)* | AIA marker |
| IK2711 | *njls63[odr-2(2b3a)p::GFP] (I)* | AIB marker |
| IK2672 | *njls39[acc-2-p::TagRFP] (I)* | AIZ marker |
| IK2951 | *njls85[sto-3p::GFP] (X)* | RIB marker |
| IK2881 | *njls83[tdc-1p::GFP] (X)* | RIM marker |
| IK3239 | *njls114[unc-47p::flp, ser-4p::frr::TagRFP] (IV)* | RIS marker |
| IK3246 | *njls119[npr-4ap::GFP] (V)* | AVA marker |
| ST401 | *ncls401[opt-3p::ArchT::GFP, acd-4p::GFP]* | AVE marker |
| IK3148 | *njls102[odr-2(18)p::GFP] (IV)* | SMBD/V marker |
| IK3147 | *njls101[lad-2p::flp, flp-22p::frr::TagRFP] (III)* | SMDD/V marker |
| IK3087 | *njls94[rig-5ap::GFP] (V)* | RMD/D/V marker |
| IK3047 | *njls88[unc-47p::GFP] (II)* | RMED/V marker |
| IK0597 | *gcy-23(nj37) gcy-8(oy44) gcy-18(nj38)* | AFD disruption |
| IK2572 | *ceh-36(ky640)* | AWC disruption |
| IK2613 | *gcy-23(nj37) gcy-8(oy44) gcy-18(nj38); ceh-36(ky640)* | AFD/AWC disruption |
| IK0604 | *eat-4(ky5)* | glutamatergic transmission disruption |
| IK0884 | *eat-4(ky5); njEx379[gcy-8p::eat-4 cDNA, ges-1p::nls-GFP]* | rescue of *eat-4* in AFD |
| IK0885 | *eat-4(ky5); njEx380[odr-3p::eat-4 cDNA, ges-1p::nls-GFP]* | rescue of *eat-4* in AWC |
| IK3331 | *njEx1387[ins-1p::flp, gcy-28dp::frr::YCX1.6]* | AIA imaging |
| IK3330 | *njEx1386[apff-1p::flp, inx-1p::frr::YCX1.6]* | AIB imaging |

## Supplemental Movie Legends

**Movie S1.** Thermotaxis behavior is accomplished within 30 minutes. Each dot represents the centroid of the animal during the thermotaxis assays in the *T<T_c_* condition (left panel) and in the *T>T_c_* condition (center panel). The time course of TTX indices in the *T<T_c_* condition (red line) and in the *T>T_c_* condition (blue line) are shown in the right panel.

**Movie S2.** Thermotaxis simulation reproduces the population behavior in the assays. Each dot represents the centroid of the animal during the thermotaxis assays in the *T<T_c_* condition (left upper panel) and in the *T>T_c_* condition (left lower panel). The animals in the thermotaxis simulation are shown in the center column. The time courses of TTX indices in the experiments (colored lines) and in the simulations (black lines) are shown in the right column.

